# Multi-Omic Dissection of Autism Reveals Dominant Effects of Family, Sex, and Host-Microbe Metabolic Interactions

**DOI:** 10.64898/2026.07.22.739782

**Authors:** Ashley G. Bell, Kiana A. West, Gary Frost, Julien Wist, Elaine Holmes, Jeremy K. Nicholson

## Abstract

Autism spectrum disorder (ASD) has been associated with gut microbiome and metabolic alterations, but reported biomarkers are inconsistent and often inadequately account for family and shared environment. We analysed 620 children, including 334 with ASD and 286 neurotypical siblings, with a median age of 6.0 years (IQR 4.0–8.0). Faecal and urine samples were collected monthly up to eight times. After quality control, 869 microbiome, 754 faecal metabolome and 787 urine metabolome samples were analysed using 16S rRNA sequencing, NMR and LC–MS metabolomics. Mixed models accounted for family, repeated sampling and biological sex. Family membership explained 46.1% of variation across microbiome and metabolome profiles, compared with 8.0% attributable to ASD. After family adjustment, ASD explained only 0.1% to 0.3% of microbiome beta diversity. No stable taxonomic biomarkers were identified, and microbial classification was poor (AUROC <0.75). Urinary metabolomics identified elevated 5-hydroxy-L-tryptophan and altered phenylalanine metabolism. Structural equation modelling found no direct associations between selected taxa and metabolites. These findings argue against a universal ASD microbiome and indicate that family context contributes more strongly than diagnosis to microbial and metabolic variation.

## Introduction

Autism spectrum disorder (ASD) is a heterogeneous neurodevelopmental condition defined by persistent differences in social communication and restricted or repetitive behaviours, including atypical sensory reactivity^1^. ASD frequently co-occurs with gastrointestinal, neurological, sleep and psychiatric conditions, which add substantial clinical and biological heterogeneity and may influence microbiome and metabolic profiles^2,3^. Genetic susceptibility and prenatal environmental exposures may also contribute to ASD aetiology^4^. However, whether observed microbiome and metabolome differences reflect ASD, co-occurring conditions or shared family environment remains unclear^3,5^.

Clinical presentation and co-occurring conditions vary across development, while biological profiles may also vary within individuals over time^2,6,7^. Studies based on single-time-point sampling may therefore obscure stable associations and combine participants with distinct biological phenotypes^3,6^. Longitudinal studies incorporating family structure, biological sex and repeated sampling, while assessing whether gastrointestinal status modifies observed associations, are needed to distinguish consistent ASD-associated signals from temporal, physiological and shared environmental variation^3,5,6^.

The prevalence of ASD has risen, where in the United States, prevalence among 8-year-old children increased from 1 in 150 (0.66%) in 2002 to 1 in 88 (1.14%) in 2008, representing a 78% rise over six years, with similar upward trends reported across Europe and Australia during the same period^8–12^. Improved awareness, expanded diagnostic criteria, and greater access to services have contributed to this increase^8,13–15^. However, ASD’s aetiology is complex and multifactorial as both genetic predisposition and a variety of environmental risk factors are known to play roles^4,16,17^. Still, no single genetic or environmental factor explains more than a subset of ASD cases, and the causes in many individuals remain unidentified^18,19^. This uncertainty in aetiology, combined with the spectrum’s breadth, highlights the need for identifying biological markers and mechanisms underlying ASD’s diverse presentations.

ASD is also commonly accompanied by co-occurring medical and psychiatric conditions^2,20,21^. Anxiety disorders, epilepsy, and gastrointestinal (GI) disturbances are particularly common comorbidities in individuals with ASD ^22^. Notably, GI problems (such as chronic constipation, diarrhoea, and abdominal pain) are among the most common non-neurological comorbidities in autism spectrum disorder (ASD), occurring up to four times more often than in neurotypical peersl^22–24^.

The presence of GI dysfunction has been associated with greater repetitive behaviours, anxiety, irritability and sleep disturbances, although associations with social communication differences are inconsistent ^22,25,26^. Studies comparing autistic individuals with and without GI symptoms have reported differences in behavioural profiles and, in some cohorts, intestinal permeability, immune activity, microbial composition and microbial–host metabolism^26^. However, these findings remain heterogeneous and do not establish ASD with GI dysfunction as a distinct biological subgroup. GI symptoms, dietary selectivity, medication use and altered gut transit can each influence microbiome and metabolome profiles independently of ASD diagnosis^3^. These observations are consistent with bidirectional communication along the gut– brain axis but do not establish microbial changes as a cause of ASD or GI symptoms^26,27^. Stratifying participants by GI status may therefore improve biological interpretation in omics studies. In this study, GI status was examined to determine whether it was associated with the observed microbial and metabolic profiles.

Building on this gut-brain axis paradigm, many studies have examined the gut microbiome in children with ASD^28–32^. While research has consistently observed differences in the gut microbiome composition between individuals with ASD and neurotypical individuals, a definitive causal link has not been established ^33–35^. Reports frequently highlight reduced overall microbial diversity in the ASD microbiome and shifts in the relative abundance of specific bacterial taxa, such as increased *Clostridium* and decreased *Bifidobacterium* ^31,36–38^. However, results across studies have been inconsistent and often difficult to reproduce^35^. Independent case-control analyses frequently identify different “dysbiotic” microbes, without a consensus microbial signature of ASD. This lack of agreement is likely due to multiple confounding factors. The ASD population is highly heterogeneous (genetically and clinically), and studies vary in participants’ ages, diets, and geographic or ethnic backgrounds, all of which substantially influence the gut microbiome^39,40^. Indeed, recent systematic reviews and meta-analyses have found little overlap in specific microbial taxa associated with ASD across different cohorts^40^. Many earlier studies have also been limited by small sample sizes, with often fewer than 100 individuals, reducing statistical power to detect true associations amid high inter-individual variability^35,40^. These knowledge gaps highlight the need for larger, more diverse, and carefully controlled investigations to clarify if there are robust microbiome and metabolite profiles linked to ASD.

This study was designed to address these knowledge gaps by leveraging a large-scale, multi-site cohort and a multi-omics, systems biology approach. We analysed urine and faecal samples from 620 children, including those diagnosed with ASD and their neurotypical siblings, recruited from two distinct regions (the United States and Saudi Arabia). By including sibling controls, the study controls for some shared genetic background and household environmental factors, while the multi-site design captures a broader range of dietary and cultural influences than single-location studies. We profiled the gut ecosystem in these participants using faecal microbiota composition characterised using 16S rRNA gene sequencing, and metabolic phenotypes were captured through untargeted analysis of both faecal and urinary samples using nuclear magnetic resonance (NMR) spectroscopy and liquid chromatography-mass spectrometry (LC–MS). This integrative strategy aims to link microbial community structure with the functional capacity of microbial and host metabolism.

Our overall aim is to discover novel microbial and metabolic biomarkers associated with ASD and to advance the understanding of biological pathways that may drive ASD heterogeneity. We hypothesised that children with ASD would harbour distinct gut microbiome configurations and metabolite profiles compared to neurotypical siblings, and that these differences could provide insights into the gut–brain biochemical interactions underlying ASD. Specifically, we sought to (i) identify taxonomic and metabolic features in faecal or urinary profiles that differentiate ASD from neurotypical development, and (ii) determine whether such features relate to clinical characteristics, potentially linking mechanistic links between gut physiology and neurodevelopment. By examining a large and diverse cohort with standardised multi-omics methods, these findings are expected to contribute new biomarkers for ASD detection and inform future research on targeted interventions, through improving our understanding and management of ASD’s complex and varied manifestations.

## Results

### Cohort and data overview

A total of 620 participants were recruited. The US cohort was recruited through 74 clinical offices representing 13 states, while an additional 65 samples were collected from King Abdullah University of Science and Technology (KAUST) in Saudi Arabia. The combined cohort comprised 334 individuals with ASD and 286 neurotypical siblings or unrelated children controls (Figure 1). Participants averaged 6.0 years old (median; ± 2.9 SD) and represented 41 ethnic backgrounds. The most common were White/Caucasian (48.1%), Latino/Hispanic (21.9%), and Asian (15.3%). The ASD group exhibited a male-to-female ratio of 2.26:1, with common comorbidities including neurodevelopmental delays (15.27%), ADHD (9.28%), and digestive issues (11.37% versus 2.09% in the neurotypical group).

**Figure 1.**
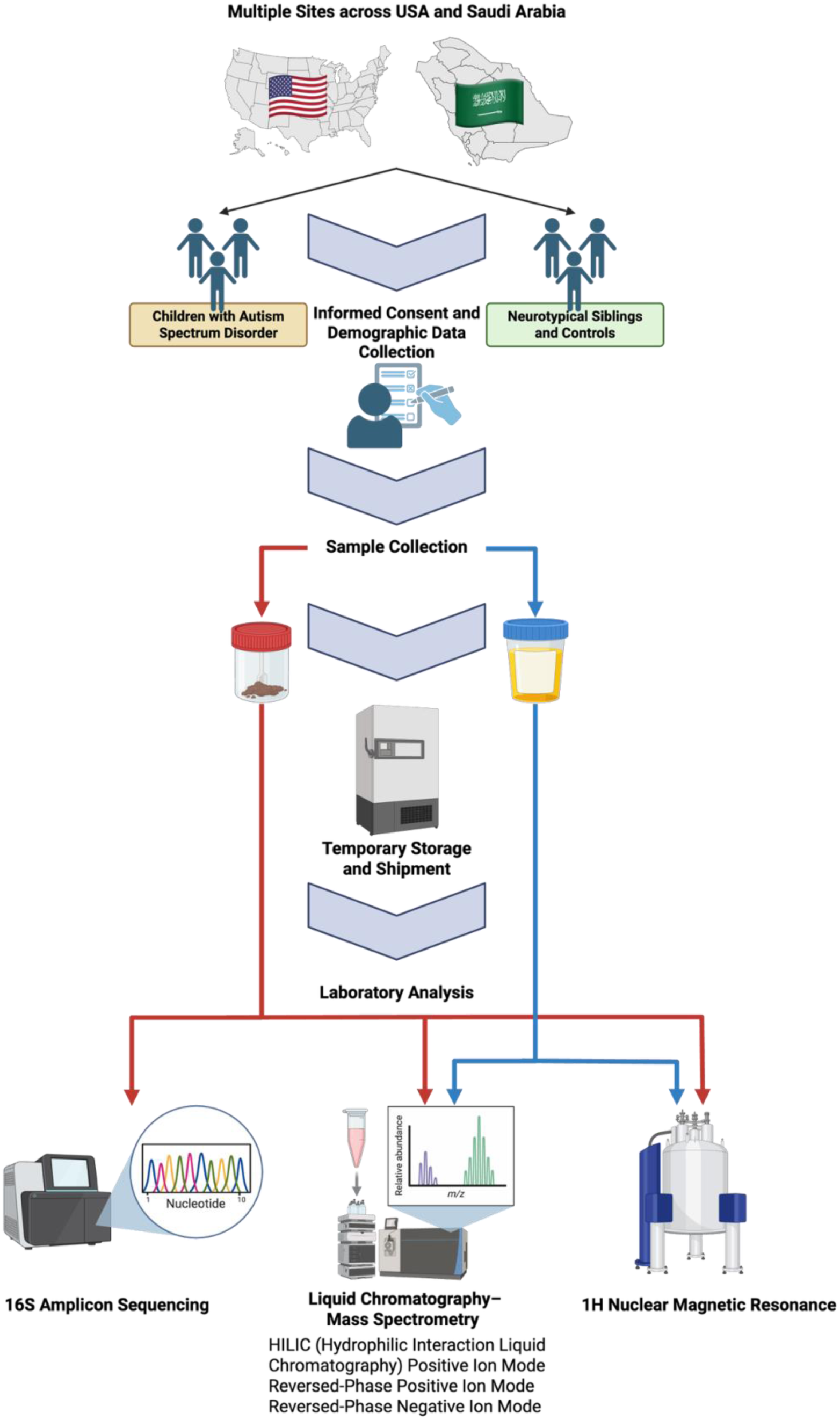
Children were recruited through CARD (USA) and KAUST (Saudi Arabia). Caregivers collected faecal and urine samples at baseline, with optional monthly follow-up for up to 8 months. Samples were shipped to Imperial College London. Faecal extracts and urine were profiled by 1H NMR and LC–MS, and faecal samples underwent 1cS rRNA gene sequencing for microbiome analysis. Image made using BioRender.

Individuals contributed up to eight samples at monthly intervals following baseline screening. After quality control, 503 urine, 430 faecal, and 503 microbiome datasets were available for analysis from the ASD group, representing 293, 267 and 284 unique individuals from each assay type, respectively. For neurotypical individuals, 284 urine, 324 faecal, and 366 microbiome datasets were available, representing 261, 235 and 253 unique individuals from each assay type, respectively. Additional metadata included dietary information, supplementation, medication, recent illness, mode of delivery, allergies, weight, height and feeding method at birth were recorded. Study design, exclusion criteria, and the ASD diagnostic method can be found in the methods section.

### Differences in familial groups overshadow differences in ASD from neurotypical sibling controls

Preliminary analyses of the uncorrected microbiome and metabolomic profiles revealed a strong family effect. Samples from the same family were more similar to each other than to samples from unrelated individuals, regardless of ASD status (Supplementary Figure 1). This is to be expected, as siblings share genetics, as well as commonalities in diet and environment, all of which are known to influence the microbiome and metabolome. This was statistically supported by PERMANOVA (Permutational Multivariate Analysis of Variance) on Euclidean distances, where family and individual identity explained substantially more variation than ASD status (Supplementary Table 1). This finding supported the need to account for family structure and repeated sampling before testing ASD-associated differences.

To reduce this confounding effect, each microbial and metabolic feature was residualised using a mixed effects model with random intercepts for family and individual nested within family. By using the residuals from the mixed effects model, the variation attributable to the family effect was removed from the dataset (Supplementary Figure 2). This allowed for a more accurate and precise comparison of the metabolic and microbial profiles between the ASD and neurotypical groups, providing a clearer understanding of the biological differences associated with the condition.

### Poor discrimination of ASD samples using metabolic and microbiota datasets

Orthogonal Partial Least Squares Discriminant Analysis (OPLS-DA) was used to compare the residualised compositional profiles between the ASD and neurotypical controls (Figure 2). Models were fitted and scaled separately for each data layer, including the faecal metabolome, urinary metabolome and gut microbiome. The OPLS-DA model was unable to effectively discriminate between individuals with ASD and the neurotypical controls. This lack of robust separation was statistically confirmed by the model’s low Q²Y score, which is indicative of poor predictive ability, and was further evidenced by the large difference between the R²Y (goodness-of-fit) and Q²Y scores, suggesting that the model was overfitting and lacked discriminatory power. Despite the overall lack of robust separation, the OPLS-DA model highlighted several features that contributed to differences between the groups. These included phenylalanine, a feature putatively annotated as phenylacetylglycine, and the genus *Bifidobacterium,* which tended to be lower in the ASD group. As phenylacetylglycine is predominantly a rodent metabolite and can be misassigned as the human metabolite phenylacetylglutamine, this annotation was considered tentative^41^.

**Figure 2.**
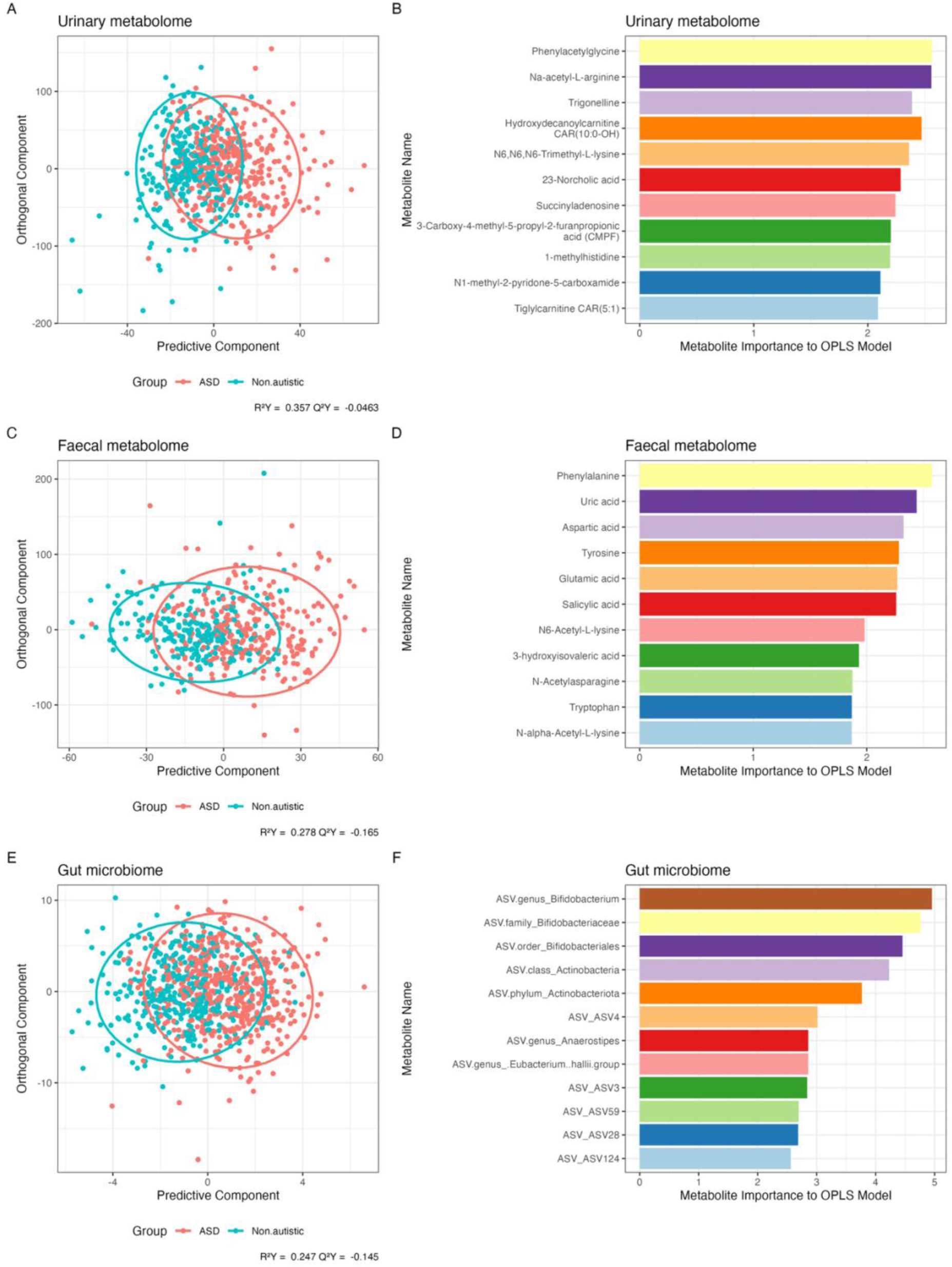
Orthogonal Partial Least Squares Discriminant Analysis (OPLS-DA) visualising key features responsible for group separation between ASD and neurotypical controls in the urinary metabolome, faecal metabolome, and gut microbiome. Panels A, C, and E display the OPLS-DA score plots for the urinary metabolites, faecal metabolites, and microbiota, respectively. These plots show the overall separation of the ASD and neurotypical groups based on their metabolic or microbial profiles. Panels B, D and F show the top-ranked discriminatory features from the OPLS-DA models, ordered by their influence on the separation between the groups.

### Most differentially abundant bacterial genera show lower relative abundance in children with ASD

Using family and individual residualised microbiome data, alpha diversity analysis showed a small but statistically significant reduction in diversity in children with ASD at the Amplicon Sequence Variant (ASV) level (Supplementary Figure 3, Supplementary Table 2). However, this difference was not observed when the diversity was aggregated at the genus level, showing that specific strains or species may be more exclusive to ASD subtypes or other environmental groupings (Figure 3). Beta diversity analysis of the residualised microbiome profiles showed a statistically detectable difference between ASD and neurotypical groups, but the effect size was negligible (Supplementary Table 3). This indicates that ASD diagnosis explained only a very small proportion of the total variance in community composition (R² values of 0.1-0.3%), suggesting no robust difference from neurotypical individuals. Despite the lack of an overall shift in community composition, differential abundance analysis at the residualised genus level data revealed that more subtle differences observed were primarily correlated with a significant decrease in the abundance of all bacterial genera in the ASD cohort. The highest losses were observed in *Desulfovibrio*, *Bifidobacterium*, and *Enterococcus* genera (Supplementary Figure 4).

**Figure 3.**
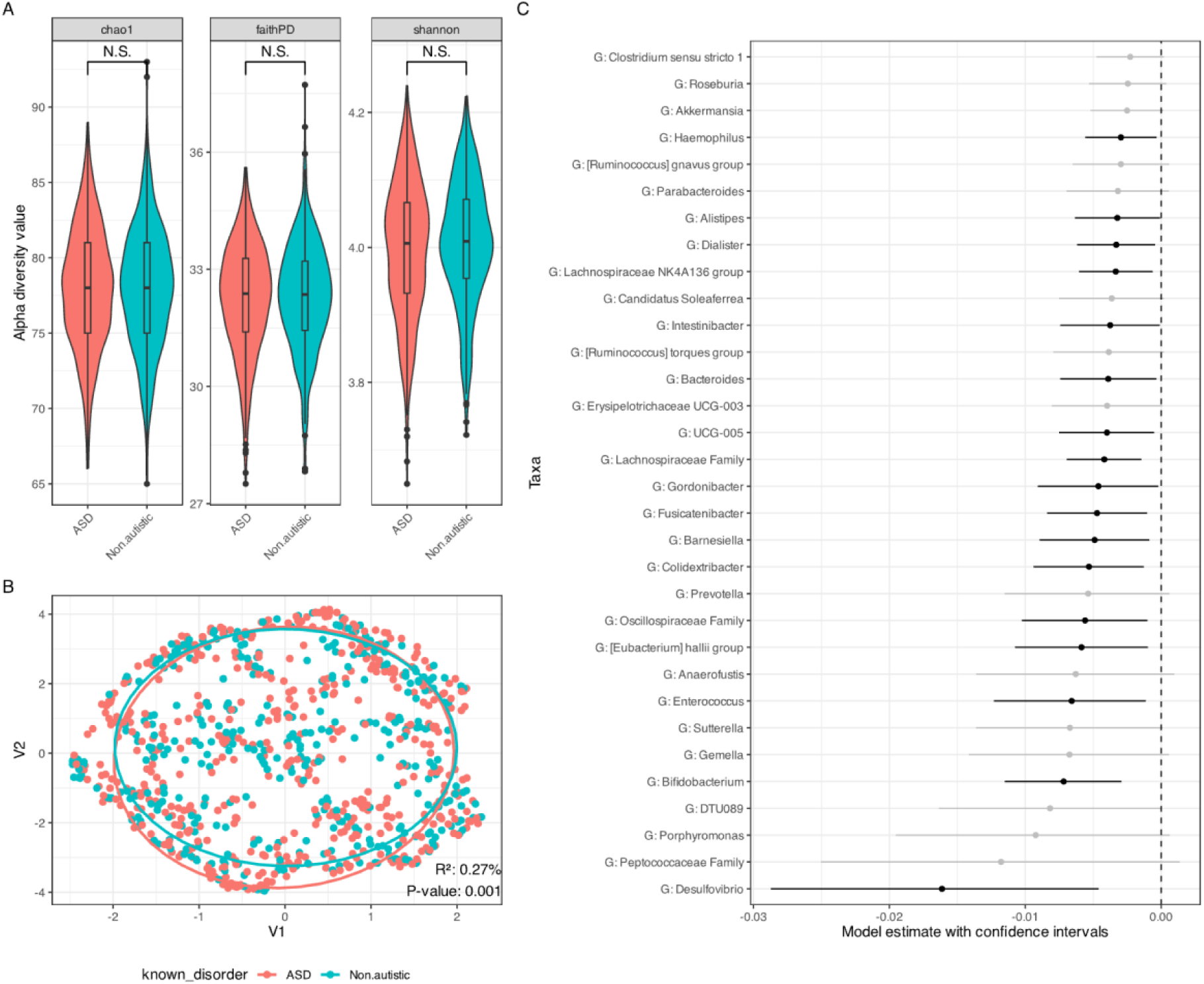
Genera Microbiome Diversity and Differential Abundance Analysis. *(A)* Alpha diversity (within-sample diversity) of the gut microbiome. Diversity was measured using three metrics: Chao1 (estimating community richness), Faith’s Phylogenetic Diversity (measuring the phylogenetic diversity of taxa), and the Shannon index (quantifying both richness and evenness). *(B)* Beta diversity (between-sample diversity) to show the overall differences in microbial community composition between the groups. The plot is based on Weighted UniFrac, a phylogenetic distance metric that accounts for both the presence and abundance of microbial taxa. *(C)* Differential abundance of bacterial taxa between the ASD and neurotypical groups, estimated using a linear model. Points represent model coefficients, and horizontal lines represent 95% confidence intervals. Negative coefficients indicate higher relative abundance in the ASD group, whereas positive coefficients indicate higher relative abundance in the neurotypical group. Black lines indicate taxa that remained significant after Benjamini– Hochberg false discovery rate correction (FDR-adjusted P < 0.05); grey lines indicate taxa that did not meet this threshold.

#### Urinary metabolites are as effective as microbiome composition or faecal metabolites at discriminating ASD from non-ASD samples

More advanced machine learning classification models (xgboost, random forest, K-nearest neighbour, native bayes, multilayer perceptron, support vector machine and logistic regression) were assessed for their effectiveness at discriminating between ASD and neurotypical samples (Figure 4). We evaluated the models based on four different input datasets: all faecal and urinary metabolites, along with microbiome taxa (green); microbiome taxa only (red); urinary metabolites only (purple); microbiome taxa and faecal metabolites (blue). The models’ performance was measured using Area Under the Receiver Operating Characteristic (AUROC) scores, where a higher score indicates better discriminatory power. The models showed similar levels of effectiveness, with the most successful approach being the model that incorporated both urinary and faecal metabolites and microbiome datasets. Although there was a clear ranking, the AUROC scores for all models were very similar, indicating that no single model provided a significantly superior discriminatory ability. This suggests that while machine learning models can find subtle differences, there is not a single, robust set of biomarkers to effectively separate the groups.

**Figure 4.**
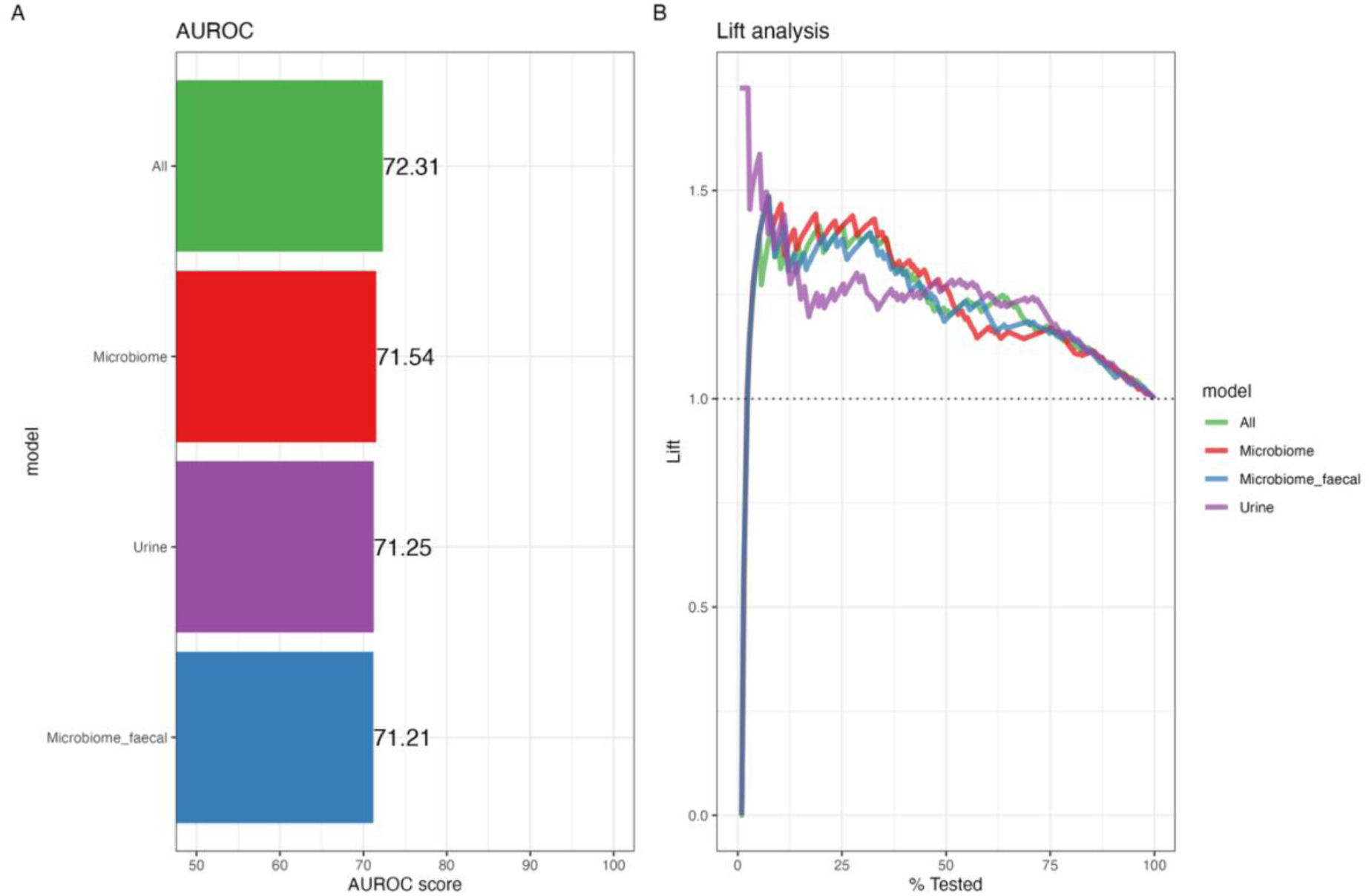
A) Discriminatory performance of different ensemble models, ranked according to their Area Under the Receiver Operating Characteristic (AUROC) score. Each model was trained on a different combination of the omics datasets to assess which dataset, or combination of datasets, was most effective at discriminating between ASD and neurotypical samples. The model was trained on: faecal and urinary metabolites and microbiome taxa (green); microbiome taxa only (red); urinary metabolites only (purple); microbiome taxa and faecal metabolites (blue). The AUROC with a score of 1.0 indicates perfect classification, and a score of 0.5 indicates performance no better than random chance. B) Lift analysis of model performance across datasets. Lift curves were generated by ranking samples according to each model’s predicted probability of ASD and calculating the cumulative proportion of true ASD cases captured as confidence decreased. A random-chance classifier is included as a dotted line at 1.0 to provide a baseline for comparison. Each curve represents a separate model trained on one dataset type (microbiome, faecal metabolites and microbiome, urinary metabolites, or combined data). The plot illustrates how effectively each model classifies ASD cases within the higher-confidence segments of its ranked predictions.

To further assess the models’ effectiveness, we analysed their performance using a lift analysis by aligning samples based on the model’s confidence in its classification as ASD or non-ASD (Figure 4). All models consistently performed better than a random-chance classifier (represented by a dotted line at 1.0), indicating a level of discriminatory power in each dataset. The model trained on urinary metabolites (purple) demonstrated the best performance. The urinary model showed high confidence in its classification of a subset of samples, and these high-confidence classifications were consistently correct. This suggests that certain urinary metabolite profiles are highly specific to ASD and that the model was able to identify a distinct subgroup with these characteristic profiles. In contrast, the other models (microbiome, faecal metabolites, and combined datasets) performed less effectively at the highest confidence levels. These models produced a higher rate of confident but incorrect classifications (false positives). This indicates that the features these models identified as being highly representative of ASD were not consistently predictive, likely due to the high biological heterogeneity within the ASD cohort. There is no single, uniform "ASD signature" in these datasets from which the models can consistently and reliably learn.

#### Extreme boosted trees and random forest models are best at classifying ASD metabolite and microbiome taxa

The contribution of different classification algorithms was assessed within an ensemble model to determine which types of models provided the most predictive power (Supplementary Figure 4). The ensemble combined Extreme Gradient Boosting (XGB), Random Forest (RF), Multi-Layer Perceptron (MLP), Support Vector Machine (SVM), Naive Bayes (NB), k-nearest neighbours (KNN), and Logistic Regression (LR). These algorithms identify group differences in different ways. Logistic regression detects relatively simple patterns in which changes across features combine additively, whereas tree-based models such as XGB and RF can detect thresholds and interactions between features. For most datasets, XGB and RF provided most of the classification power, suggesting that separation between the ASD and neurotypical groups depended mainly on combinations of microbiome and metabolic features. In the urinary metabolite ensemble, however, LR, MLP and SVM also contributed substantially. This indicates that part of the urinary signal could be captured through relatively simple, coordinated differences across metabolites, while other parts required more flexible combinations of features. Combining these models therefore improved classification because each captured a different aspect of the urinary data.

### ASD is characterised by *Bifidobacterium* and *Holdermanella*, and urinary metabolites such as tryptophan

SHapley Additive exPlanation (SHAP) values were used to assess the importance and direction of features contributing to each ensemble model (Figure 5). n models incorporating microbiome data, the direction of association varied considerably between models, indicating substantial inter-individual heterogeneity. For example, Bifidobacteriaceae was associated with ASD in the model combining all datasets but with neurotypical individuals in the faecal metabolite and microbiome model. Holdemanella was also an important feature in several models, but its direction of association was inconsistent (Supplementary Figure 5). Although Actinomycetes and Porphyromonadaceae were consistently associated with neurotypical individuals, the overall variability provided little evidence for robust microbial classifiers.

**Figure 5.**
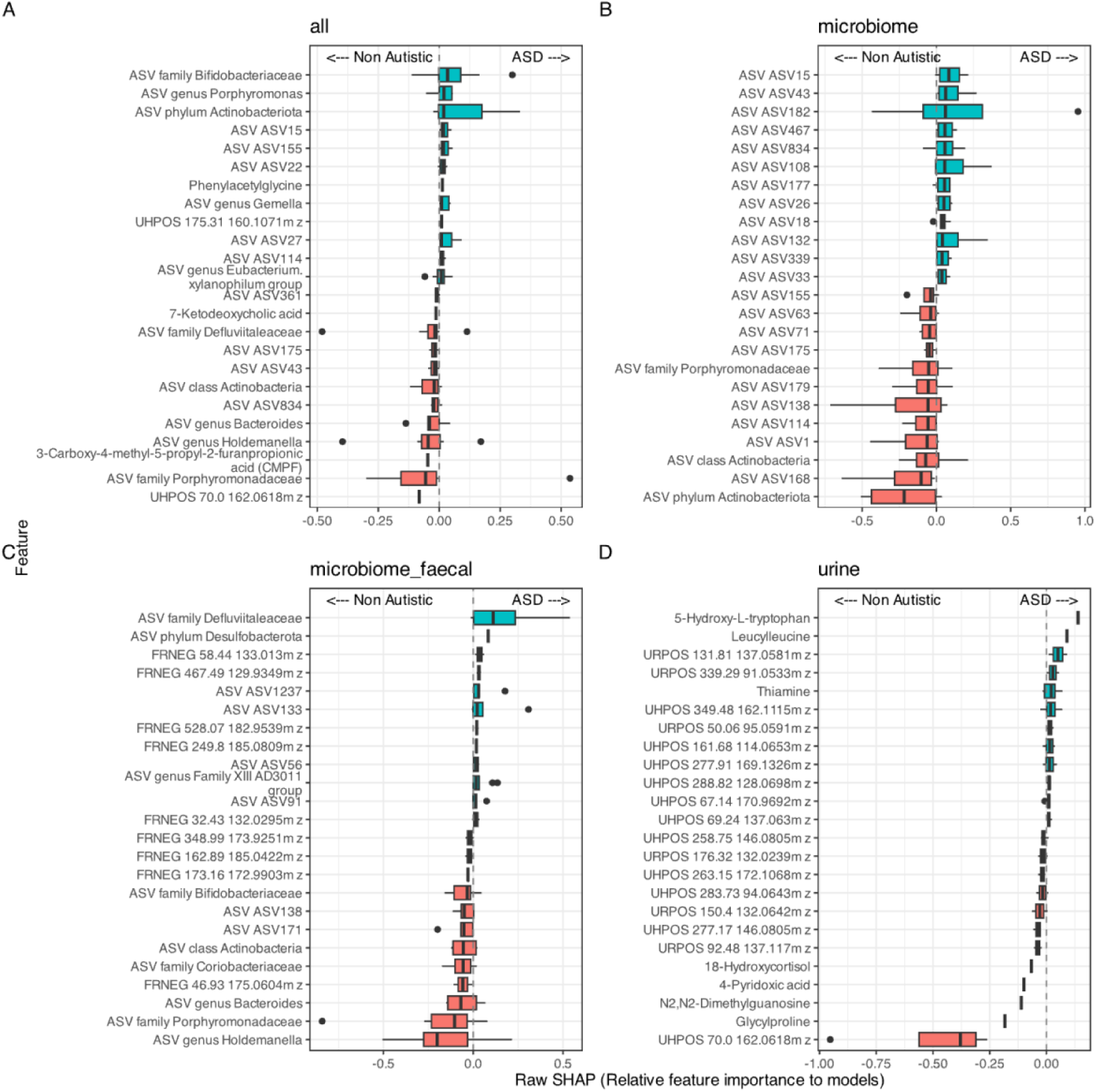
SHapley Additive exPlanations (SHAP) Scores of Ensemble Models representing SHAP analysis of the ensemble models to explain the contribution of top features to the models’ predictive power. Each boxplot represents a feature, with its position on the x-axis indicating the feature’s influence on the model’s prediction (positive values for ASD classification, negative for neurotypical classification). (A): All datasets, including faecal and urinary metabolites and microbiome taxa. (B): Microbiome taxa only. (C): Urinary metabolites only. (D): Microbiome taxa and faecal metabolites.

The urinary metabolite model presented a different pattern, with its top features showing a much clearer and more consistent directionality. However, in the model trained with only urinary metabolites, 5-Hydroxy-L-tryptophan and Leucylleucine were almost always associated with the ASD phenotype, whereas Glycylproline, N2, N2-Dimethylguanosine, 4-Pyridoxic acid, 18-Hydroxycortisol and unidentified metabolite UHPOS 70.0 162.0618m/z were associated with the neurotypical group. This suggests that the urinary metabolome may contain more specific markers for both ASD and neurotypical profiles. While some faecal metabolites were found to be associated with ASD and neurotypical phenotypes, we could not reliably determine their identity.

#### Biological sex likely influences biomarkers of ASD

Biological sex was identified as a feature with the highest model predictive power, suggesting that the biological differences between sexes are important differentiators of the microbiome and metabolome in this cohort, with consistent results across models (Supplementary Figure 6). Several lifestyle and early-life factors were found to have minimal predictive power within the models. These included ethnicity, mode of delivery (Caesarean section vs. vaginal birth), and type of milk feeding at birth (breast, formula, or mixed). This suggests that these factors did not create a strong enough biological signature to be useful for classifying samples. However, it is important to note that this lack of predictive power may be attributed to smaller sample sizes within these specific subgroups. For instance, the number of participants born through caesarean section or who were formula-fed may have been too limited to provide a robust statistical signal, reducing models’ ability to identify a clear relationship.

#### The relationship between ASD linked bacteria and subsequent metabolite production is likely more complex

To identify causal links between metabolites and the bacterial microbiome, top bacterial genera identified through SHAP scores were correlated with top faecal metabolites identified to be strongly associated with ASD diagnosis (Figure 6). Structural Equation Modelling (SEM) was used to test for a direct causal relationship from the ensemble model of metabolites and bacterial genera that were significantly correlated with each other. The SEM did not support a direct causal relationship between any of the tested bacterial genera and metabolites, likely due to the low correlation values. This negative finding suggests that if there is a relationship between the gut microbiome and the faecal metabolome driving ASD, it is not direct, with more complex connections likely, or involving indirect pathways influenced by a variety of confounding factors, such as diet or host genetics.

**Figure 6.**
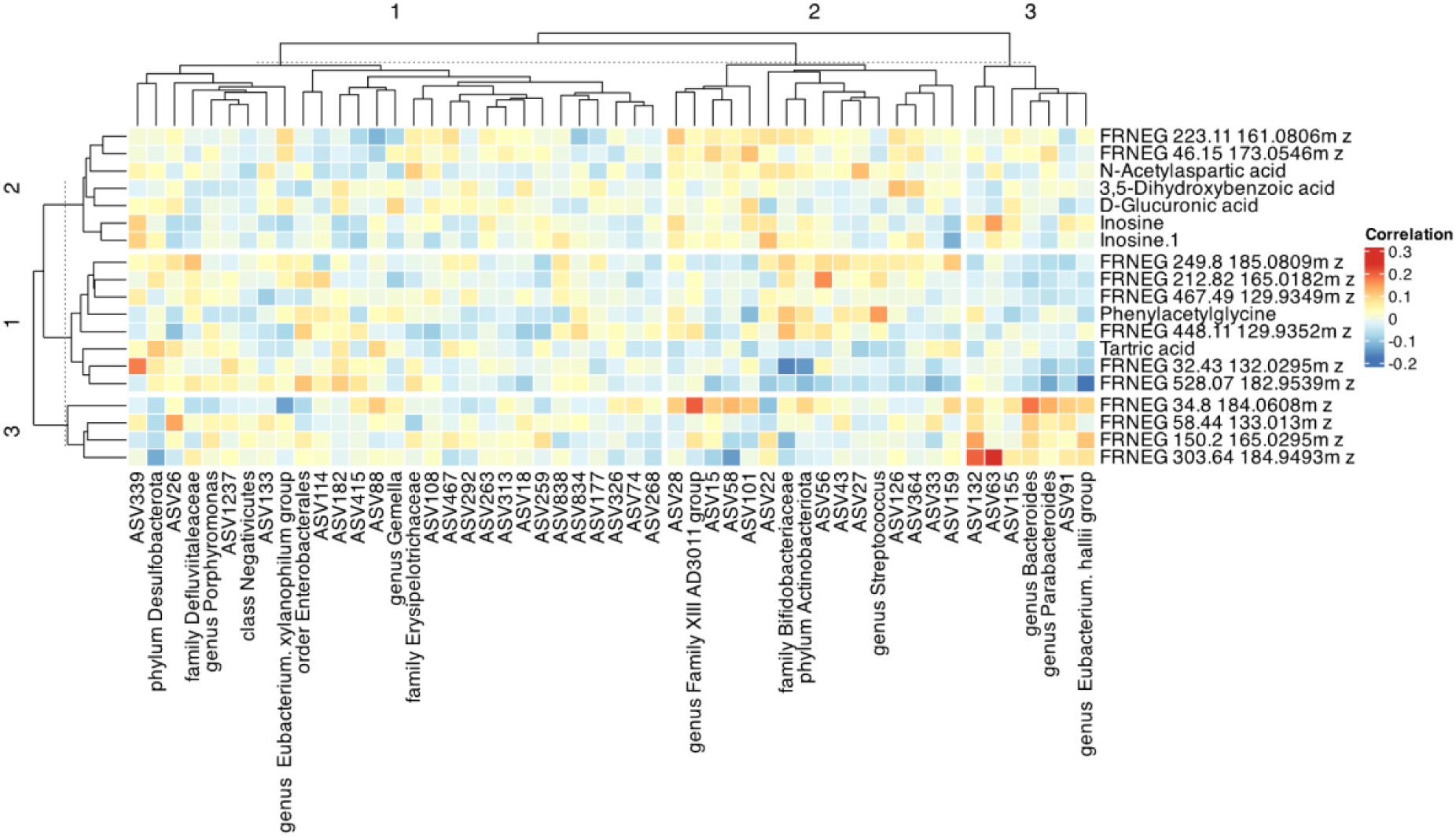
Correlation plot between metabolites and taxa with the highest model importance across ensemble models.

#### ASD sub-types are difficult to differentiate and more likely correlated with microbiome taxa compositions

Attempts to identify and cluster distinct ASD subtypes by combining biological features with specific metadata were unsuccessful. Factors such as mode of birth (e.g., Caesarean section), presence of neurodevelopmental delays indicating more severe ASD, or gut issues did not create distinct profiles in the dataset. A PERMANOVA showed metadata clusters were indistinguishable from their controls, suggesting poor discrimination of ASD subtype clusters (Supplementary Table 5). Supervised SOMs provided better classification; however, they struggled to differentiate between true and false positives, suggesting overfitting and that the measured metadata features do not create specific, well-defined subtypes in this cohort, likely due to large amounts of unfilled or missing metadata. Models with the inclusion of microbiome data seem to have better discriminatory power with higher accuracy and sensitivity, particularly when clustered by neurodevelopmental delays (Figure 7).

**Figure 7.**
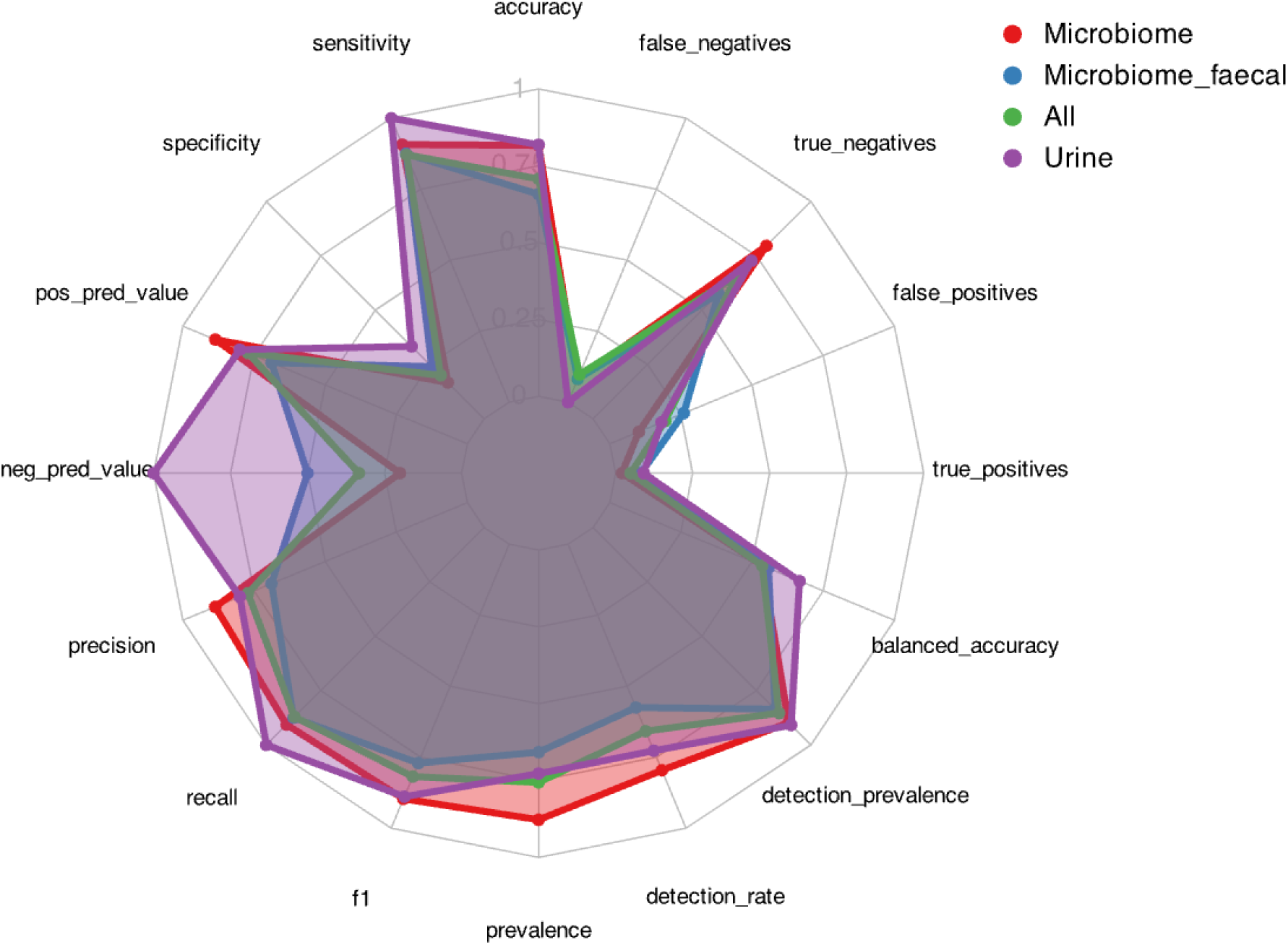
Confidence matrix of a supervised Self-Organising Maps (SOM) predicting the cluster of neurodevelopment delays using different dataset subsets.

## Discussion

Our study presents one of the largest family-matched, multi-omic investigations into the gut microbiome and metabolome in ASD to date. Using a cohort of 620 individuals and over 1,300 microbiome and metabolite profiles, we found that family-level factors accounted for the largest proportion of biological variance. Importantly, ASD status still explained 8% of total variance, indicating a measurable biological signal despite substantial family, environmental and inter-individual heterogeneity. No consistent microbiome or metabolomic signature characterised all ASD cases, but depletion of *Bifidobacterium* and disruption of serotonin-related urinary metabolites emerged as reproducible features in specific subgroups. These findings support a shift toward stratified, multi-omic models of ASD biology rather than universal biomarkers.

Family membership explained 46.1% of the total variance in microbiome and metabolome profiles, compared to 8% attributable to ASD diagnosis. This disparity supports evidence that shared environment, diet, and early-life exposures have a stronger influence on gut biology than diagnostic status^42,43^. By using neurotypical siblings as matched controls and applying a Linear mixed model, we accounted for family structure and removed shared variance. This approach provides greater specificity than case–control studies using unrelated participants, which are more susceptible to lifestyle confounding. Even after controlling for family, ASD explained only 0.1–0.3% of beta-diversity (PERMANOVA), and classification models performed poorly (AUROC < 0.75), with unstable feature directionality. These findings align with twin and sibling studies showing weak or absent microbiome or metabolomic signatures once background is matched ^44,45^.

Reported microbial taxa differences in ASD remain inconsistent across studies. Meta-analyses have found high heterogeneity in taxa-level results, with poor agreement on which bacteria are enriched or depleted in ASD cohorts ^46–48^. While taxa from the Firmicutes, Bacteroides and *Clostridium* are variably reported, effect sizes and direction are modest, inconsistent and cohort-dependent ^35^. Our results extend this by showing that within-group variability among ASD participants equals or exceeds between-group differences, further supporting the absence of a universal ASD microbial signature.

*Desulfovibrio* abundance presented conflicting results, where it was depleted in differential testing but variable in SHAP models. Prior studies implicate sulphate-reducing bacteria in ASD via hydrogen sulphide and propionate production ^49,50^. These inconsistencies likely arise from functional heterogeneity masked by 16S genus-level aggregation. Strain-level effects may drive biology despite low overall abundance, underscoring the need for metagenomic and metabolomic integration in future ASD microbiome studies.

Despite cohort heterogeneity, *Bifidobacterium* depletion was consistently observed in a large subset of ASD participants across OPLS-DA, differential abundance analysis, and SHAP scoring. As a dominant early-life commensal, *Bifidobacterium* plays a key role in fermenting dietary fibre into short-chain fatty acids (SCFAs), which supports immune regulation and gut barrier integrity, and its reduction has been reported across several ASD studies and meta-analyses ^23,51,52^. We also observed co-depletion of *Holdemanella*, another SCFA-producer, suggesting broader fermentative disruption specifically affecting butyrate biosynthesis. ASD participants had higher rates of gastrointestinal symptoms (11.37% vs 2.09%), consistent with reports linking lower faecal butyrate levels to gastrointestinal symptoms ^53,54^. Prior intervention studies demonstrate that microbiota transfer or *Bifidobacterium* supplementation can increase abundance and improve behavioural and intestinal outcomes ^53,55,56^. These findings support a model in which depletion of SCFA producers contributes to epithelial barrier dysfunction, immune activation, and altered neuroimmune signalling, which in turn suggests a multi-step gut-brain pathway rather than a direct causal metabolite effect.

Urinary metabolic models showed the most promise at classifying ASD samples compared to microbiome-based classifiers. This likely reflects the fact that urine captures functional host–microbial metabolic output with reduced sensitivity to compositional variability inherent to microbial datasets. Several studies have reported strong classification performance using urinary metabolites alone. For example, random forest models have achieved accuracies up to 85% and AUCs of 0.9 ^57^, while discriminant analysis has shown correct classification in 88.2% of ASD cases^58^. These findings raise the possibility that urinary metabolomics may offer a practical, non-invasive biomarker avenue for stratifying or screening ASD, although controlling for family remain ongoing challenges as unaffected siblings of autistic children often exhibit intermediate urinary profiles, suggesting familial (and therefore environmental) metabolic shifts rather than discrete disease-specific markers ^44^.

Among discriminatory metabolites, 5-hydroxy-L-tryptophan (5-HTP) was enriched in ASD participants, corroborating findings from other studies ^55,59^. As the immediate precursor to serotonin, 5-HTP is converted to 5-HT via aromatic L-amino acid decarboxylase ^55^. Prior studies report reduced urinary 5-HIAA/5-HTP ratios in ASD, consistent with impaired serotonin turnover, and whole-blood hyperserotonemia is observed in ∼25–30% of autistic individuals ^55,60^. This metabolic bottleneck may disrupt gut motility, neuroimmune function, and CNS development. Tryptophan metabolism is highly susceptible to microbial modulation. Gut bacteria influence both serotonin synthesis and the formation of neuroactive tryptophan derivatives such as indoleacetic acid and kynurenine ^61^. These findings exemplify indirect host–microbe interactions in which microbial metabolites modulate host enzymatic pathways with neurochemical consequences. Together, these data support a model in which ASD-associated metabolic profiles reflect systemic effects of disrupted microbial–host co-metabolism rather than isolated microbial shifts alone.

OPLS-DA analysis identified phenylalanine and its microbial-host co-metabolite phenylacetylglycine as discriminatory features between ASD and control samples. However, this annotation should be interpreted cautiously, as phenylacetylglycine can be misassigned in as phenylacetylglutamine human urinary metabolomic datasets ^41^.

We therefore interpret this signal as evidence of altered phenylalanine related metabolism, rather than as a confirmed change in phenylacetylglycine/phenylacetylglutamine specifically. Phenylalanine is an aromatic amino acid that is metabolised by phenylalanine hydroxylase into tyrosine ^62^. In classical phenylketonuria (PKU), mutations in the PAH gene reduce enzymatic activity, causing elevated phenylalanine levels and resulting in neurodevelopmental impairment and behavioural traits resembling autism ^63^. While it is unknown if any participants in this study met diagnostic criteria for PKU, elevated urinary phenylalanine suggest a partial disruption of aromatic amino acid metabolism in a subset of ASD individuals. Phenylacetylglutamine is produced when excess phenylalanine is conjugated with glutamine in a detoxification pathway, and its increase further supports dysregulated phenylalanine handling ^62^. Similar patterns have been reported in previous ASD metabolomic studies, which observed higher urinary phenylalanine concentrations compared to controls and suggested altered PAH activity or microbial processing of phenylalanine ^59,64,65^. These findings support the potential of urinary metabolomics to identify host metabolic deviations not captured by microbial or genomic profiling alone.

To test whether differences in microbial composition directly explain shifts in faecal metabolite levels, Structural Equation Modelling (SEM) was applied to evaluate causal relationships between the top differentially abundant bacterial genera and their most strongly correlated metabolites. The analysis identified no statistically supported direct linear paths, consistent with low correlation coefficients observed across taxa metabolite pairs. This negative finding suggests that microbial composition alone does not explain metabolite differences in ASD. This may reflect indirect host mediated effects, but also functional redundancy within the microbiome, where different bacterial taxa can perform similar metabolic roles. As a result, the same metabolite profile may arise from different community structures, weakening simple taxon metabolite correlations. These findings support an indirect and functionally distributed model of host microbe interaction, where metabolic outcomes are shaped by both host processes and overlapping microbial functions. These include gut epithelial barrier integrity, mucosal immune response, hepatic metabolism of microbial products, and systemic hormonal signalling pathways. Similar conclusions have been drawn from prior microbiome–metabolome studies showing that variation in gut microbiota composition explains only a minority of variation in metabolite profiles, especially when host absorption, transport, and enzymatic transformation are involved^66–68^ This highlights the limitations of compositional microbiome data for inferring metabolic function and supports the need for integrated, multi-layered models when interpreting gut–brain axis effects in ASD.

Biological sex was the most consistent predictive variable across all microbiome, metabolome, and combined models. This result aligns with the known male predominance in ASD diagnosis, reflected in our cohort ratio of 2.26:1. However, the high feature importance of sex does not imply that one sex is intrinsically more predictable. Instead, it suggests that different biological processes may underpin ASD in males and females. Emerging data support sex-dependent variation in microbiome and metabolome profiles within ASD populations. For example, studies report distinct microbiota–epitope interactions in males with ASD but not in females, indicating sex-specific immune–microbiome relationships ^69^. Similarly, microbiome composition has been shown to differ by sex among autistic individuals, further supporting differential biological architecture ^70,71^. These findings highlight that ASD is not a biologically uniform condition and support the need for interaction-based modelling approaches. Importantly, sex should not be treated solely as a nuisance covariate but as a biological axis potentially moderating both risk and presentation. Given the unbalanced sex distribution in this cohort and the limited power to resolve within-sex subtypes, these findings should be interpreted as hypothesis-generating rather than definitive.

Taken together, these findings argue against the existence of a single, reproducible microbial or metabolic marker for ASD. Rather than a direct, linear pathway linking specific bacterial taxa to metabolite changes and behavioural phenotypes, the evidence supports a model in which gut-derived signals are processed through systemic host mechanisms. These include hepatic metabolite transformation, immune signalling, endocrine modulation, and barrier integrity, each of which may act as a mediator between microbiome composition and brain function. The demonstrated importance of family, biological sex, and urinary metabolomic patterns further highlights the need for a stratified research framework. Future work should prioritise longitudinal, family-based or twin cohorts to track developmental trajectories; integrate metagenomic and metabolomic profiling across gut, plasma, and urine compartments; and explicitly model host–microbe interactions using systemic markers such as cytokines or neurotransmitter precursors. Stratification by sex and identification of biologically coherent subgroups—such as individuals with disrupted SCFA metabolism or altered serotonin turnover—will be essential for developing clinically meaningful diagnostics or interventions. The concept of a universal “autism microbiome” is not supported by these data. Instead, precision models incorporating host–microbe co-metabolism, genetic background, and individual-level features will be required to realise the translational potential of gut–brain axis research in ASD.

## Methods

### Study population and sampling

Children in the United States were recruited through Center for Autism and Related Disorders (CARD) offices between December 2015 and June 2017.

Individuals were asked to donate a faecal and a urine sample upon entering the study, with the option of donating additional urine and faecal samples monthly for up to 8 months (Figure 8). Biospecimens were collected by the caregiver, stored at -20°C upon collection and kept frozen until being stored at -80°C within a CARD centre. Samples were shipped to Imperial College London on dry ice. For each collection time point, caregivers were asked to complete a questionnaire detailing the child’s age, height, weight, current medical issues, medications and dietary information. ASD diagnosis was performed using the DSM IV and V. There were no exclusion criteria.

**Figure 8.**
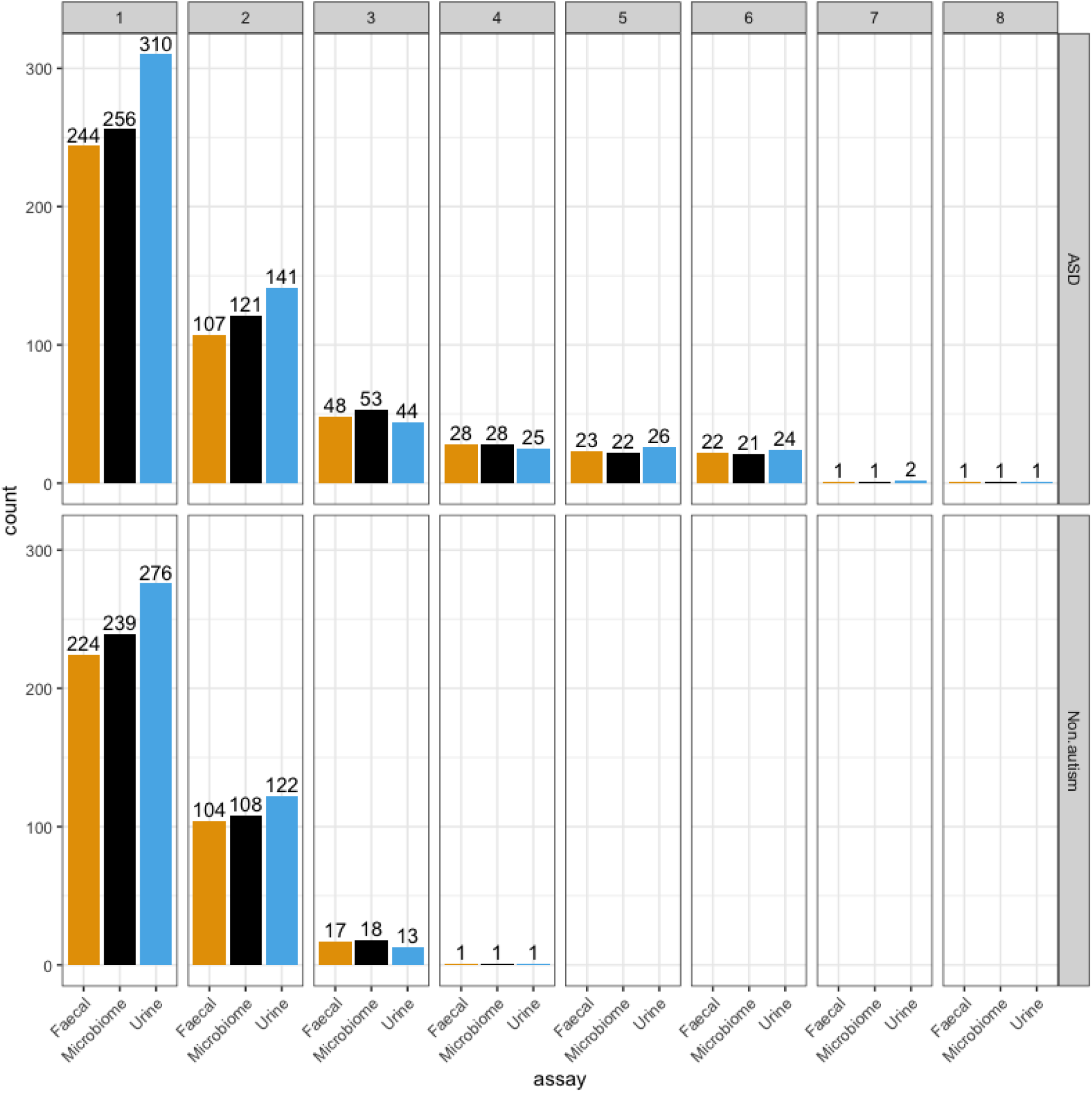
Total counts of samples split by ASD status for each timepoint. Availability of each sample labelled in colour with microbiome representing 16S amplicon and urine and faecal representing metabolic

### Metabolic extraction

Metabolites were extracted from faecal samples using acetonitrile buffer (ACN):H2O (1:3) containing 2% w/v sodium azide and bead-beating. Tubes containing 625 μl buffer, 250 mg faeces and 1.0 mm Zirconia beads (Stratech) were shaken for 45 sec in a Biospec bead-beater and centrifuged for 20 min at 17,000 x g and 4°C. Supernatant was filtered through 0.45 μm Spin-X® centrifuge tube filters (Corning) and stored at -80°C. Global metabolic profiles of faecal extracts and urine samples were acquired by 1H NMR spectroscopy and LC-MS using HPOS, RPOS and RNEG techniques at the Imperial Phenome Centre (Imperial College London, UK). For NMR samples A standard one-dimensional pulse sequence with water presaturation was used.

### DNA Extraction

DNA was extracted from 250 mg of each sample using the PowerLyzer PowerSoil DNA Isolation Kit (Mo Bio, Carlsbad, CA, USA). Samples were homogenised by bead beating in a Bullet Blender Storm (Chembio Ltd., St. Albans, UK) for 3 min at speed 8 to ensure cell lysis. The Qubit fluorometric assay (Thermo Fisher Scientific, Carlsbad, CA, USA) was used to quantify DNA. The V1-V2 region of the 16S rRNA gene was amplified by PCR using a mixture of 4 forward primers at 4:1:1:1 (28F-YM:28F-Borrellia:28FChloroflex:28F-Bifdo) and 1 reverse primer (388R). SequalPrep Normalisation Plate Kits (Life Technologies, Paisley, UK) were used to clean and normalise PCR reactions, which were then pooled into libraries. Libraries were quantified using the NEBNext Library Quant Kit for Illumina (New England Biolabs, Hitchin, UK) prior to being sequenced.

### Metabolic processing

All NMR metabolomic data from urine and faecal samples were processed using the nPYc toolbox (v2.0). Raw Bruker files were imported following the standard nPYc one-dimensional proton NMR pipelines (“noesygppr1d” acquisition) using the GenericNMRurine SOP. Spectra underwent automated processing within nPYc, including Fourier transformation, phasing, baseline correction, chemical shift calibration, and spectral alignment according to the default workflow. Samples failing automated quality control criteria were excluded based on line width, calibration accuracy, baseline quality, or solvent peak interference. Processed spectra were normalised using probabilistic quotient normalisation to account for dilution effects prior to downstream analysis.

For LC–MS datasets, samples acquired in HILIC (Hydrophilic Interaction Liquid Chromatography) Positive Ion Mode (HPOS), Reversed-Phase Positive Ion Mode (RPOS) and Reversed-Phase Negative Ion Mode (RNEG) modes for urine and faecal matrices were processed separately. Raw Bruker files were converted to mzML using msConvert (ProteoWizard) and processed in R using the Bioconductor packages xcms and CAMERA following the nPYc workflow. Chromatographic peaks were detected using the CentWave algorithm, refined by merging neighbouring peaks, grouped across samples using a density-based approach, and retention time aligned using the peakGroups method. Missing peak areas were integrated to generate a complete feature intensity matrix. Putative isotopes and adducts were annotated using CAMERA.

Feature tables were imported into nPYc for structured quality control. Features were filtered based on correlation to study reference samples (>0.7), relative standard deviation (<30%), and variance ratio (>1.1). Samples exceeding the 95th percentile for DmodX, Hotelling’s T², or scores distance were excluded. Batch correction was applied within nPYc, and data were normalised using probabilistic quotient normalisation. To obtain targeted relative concentrations for metabolites of interest within the LC–MS datasets, PeakPantheR was applied within the nPYc framework using the same parameter settings as the untargeted workflow.

### 16S rRNA gene sequencing and processing

Paired-end 16S rRNA gene amplicon sequencing data were processed in R using the DADA2 pipeline (v1.26). Forward and reverse reads were quality assessed using FastQC, and primer removal was confirmed prior to analysis. Based on read quality profiles, forward and reverse reads were truncated at 265 bp and 190 bp, respectively, to retain high-quality bases while maintaining sufficient overlap for merging. Reads containing ambiguous bases or exceeding expected error thresholds were removed (maxEE = 2), and PhiX sequences were filtered.

Error rates were learned separately for each sequencing run to account for run-specific error profiles. Amplicon sequence variants (ASVs) were inferred using the DADA2 denoising algorithm without pooling, followed by paired-end merging. A sequence table was constructed and filtered to retain ASVs within the 1st–99th percentile of the observed length distribution to remove anomalous sequences. Chimeric sequences were identified and removed using the consensus method.

Taxonomic assignment was performed using the SILVA reference database (v138.1), with classification retained to genus level. A phyloseq object was constructed integrating the ASV table, taxonomy, and sample metadata. Non-bacterial sequences (mitochondria, chloroplasts, Eukaryota) were removed.

Contaminant ASVs were identified using the decontam prevalence method with negative controls and removed prior to downstream analysis. Negative controls were excluded from subsequent analyses. Samples with substantially reduced sequencing depth were assessed but retained, as depth variation did not exceed one order of magnitude.

A phylogenetic tree was constructed from ASV sequences using MAFFT for multiple sequence alignment and IQ-TREE for maximum likelihood inference with model selection. The resulting tree was rooted using the longest terminal branch and incorporated into the phyloseq object for downstream diversity analyses.

### Data harmonisation and family-level correction

Multi-omics datasets (untargeted LC–MS, NMR, targeted LC–MS and 16S amplicon data) were assembled and harmonised in R using a unified MultiAssayExperiment object. Post-import filtering was applied to each assay to remove features with all missing values and features with >30% zero values across study samples.

For 16S assays, counts were transformed using robust centred log-ratio (rclr) prior to downstream modelling, whereas metabolomic datasets were z-scaled transformed. Features with near-zero variance were removed per assay (caret nzv). To account for relatedness and repeated sampling, each feature was residualised using a linear mixed-effects model with random intercepts for family and individual nested within family, (1 | Sibling_pair/CollectionID); if the model was singular, a reduced structure (1 | Sibling_pair) was used. Residuals were used as family-corrected values and concatenated across assays into a single analysis matrix. Curated covariates (diagnosis group, ethnicity, mode of delivery, sex, feeding at birth) were merged with the corrected data to generate the final analysis dataset.

### OPLS-DA

OPLS–DA was performed in R using ropls on the family-corrected multi-omics matrix. Separate models were fitted for (i) urine metabolites (targeted PeakPantheR and untargeted LC–MS/NMR), (ii) faecal metabolites (targeted PeakPantheR and untargeted LC–MS/NMR), and (iii) faecal microbiome features (16S abundance tables). Samples with missing values were excluded. Models were fitted with one predictive and one orthogonal component (predI = 1, orthoI = 1), standard scaling (scaleC = "standard"), 7-fold cross-validation (crossvalI = 7), and permutation testing (permI = 30). Model performance was summarised using cumulative R²Y and Q². Feature contributions were ranked using VIP scores reporting only top-ranked features.

### Microbiome analysis

Microbiome analyses were performed in R using phyloseq, vegan, microViz. Analysis was only performed on ASV- and genus-level count on pre-processed data (rclr; near-zero variance removed; family corrected) as described above. Diversity metrics were estimated from 1,000 resampling iterations per taxonomic level. For each iteration, sequences were resampled with replacement, with sample size set to 95% of the minimum library size. Alpha diversity was computed as Shannon, Chao1, and Faith’s phylogenetic diversity. Beta diversity was computed as robust Aitchison distance, Bray-Curtis, and Generalised UniFrac (α = 0.5 and α = 1; unweighted UniFrac).

Associations with ASD were tested per iteration. Alpha diversity was tested using logistic regression with ASD diagnosis as the outcome and the diversity metric as the predictor; effect estimates were summarised by the median across iterations. Beta diversity was tested using PERMANOVA (vegan::adonis2, 999 permutations) on each distance matrix; R² and P-values were summarised by the median across iterations. For visualisation, distance matrices were aggregated across iterations and embedded using UMAP (n_neighbors = 30, min_dist = 0.01).

Differential abundance was assessed per iteration using a multivariable binomial GLM including taxa as predictors, followed by backward stepwise selection. P-values were adjusted using Benjamini–Hochberg, and taxa were retained if selected in >30% of iterations; effect sizes and confidence intervals were summarised using the median across iterations.

### Model classification

Classification models were trained in R using tidymodels, with an 80/20 stratified train/test split and 10-fold stratified cross-validation on the training set. Within each recipe, transformed datasets (z-scaled metabolomics; rclr 16S) with all assay predictors missing were removed, near-zero variance predictors were dropped, linear combinations and highly correlated predictors were removed (Pearson r > 0.9), missing values were imputed using k-nearest neighbours.

Seven model classes were tuned per assay using a workflow set: random forest (ranger), gradient boosting (xgboost), RBF SVM (kernlab), kNN (kknn), naïve Bayes (klaR), multilayer perceptron (h2o), and penalised logistic regression (glmnet). Hyperparameters were tuned via grid search (20 settings) with AUROC as the optimisation metric.

Ensemble models were built using the R package stacks by blending all models for each assay with elastic-net regularisation (penalty grid 10^-3 to 10^2; mixture 0 to 0.5), repeated 30 times, and fitting member models with non-zero ensemble coefficients. Ensembles were trained for four feature sets: microbiome-only, microbiome plus faecal metabolomics, urine-only, and all modalities. Final performance was evaluated on the held-out test set using AUROC, ROC curves, and lift curves.

Model interpretation used SHAP values computed per contributing ensemble member. Member models with positive ensemble coefficients were extracted and SHAP values were estimated using fastshap::explain (nsim = 30). For each ensemble model, mean absolute SHAP and mean signed SHAP were calculated per feature and scaled within each model. Model-level SHAP summaries were combined by weighting feature SHAP values by the ensemble coefficient, then rescaled within each ensemble to compare feature importance across experiments. Top features were reported using signed SHAP (direction towards ASD vs non-autistic) and absolute SHAP (overall importance).

### Structural Equation Modelling analysis

For correlation analysis, the top 24 ASD-associated features (ranked by median signed SHAP within each experiment) were selected. The corresponding z-scored transformed feature columns were pairwise Pearson correlated. A microbiome–metabolite correlation matrix was then formed by retaining metabolite rows and ASV columns, and visualised as a clustered heatmap (k-means row/column splitting, k = 3).

A structural equation model (SEM) was fit in lavaan to test mediation structures by linking microbiome features, metabolite features and subsequent ASD status. Latent variables were defined as a microbiome component and a metabolite profile. Directed paths were specified from microbiome component to metabolite profile, and from metabolite profile to ASD status, with standardised estimates reported from the fitted SEM.

### Self-Organising Maps clustering

ASD-only samples were subsetted with separate analysis tables for urine features, microbiome-only features, faecal metabolites plus microbiome features, and the full feature set.

To test whether metadata variables explained overall ASD feature structure, a supervised self-organising map (SOM) was fit using kohonen::xyf to predict a binary clinical label (e.g. neurodevelopment delays). Data were split into 75% training and 25% testing splits. Model performance was evaluated on the held-out test set using the predicted class. Confusion-matrix derived statistics (e.g. true/false positives) were reported and visualised as a radar chart.

## Declarations

### Ethics approval

The study was approved by the CARD Institutional Review Board (IRB00004971), and written informed consent was obtained from a legal guardian of each participant. Separately, children were recruited in Saudi Arabia through during the same period. The same sampling and storage procedures were used for both cohorts.

### Data availability

Full processing parameters and reproducible code are available in the project repository https://github.com/ash-bell/jebsen_CARD. 16S amplicon sequences can be found under the EBI/ENA accession number PRJEB121195. Metabolomic raw files can be found under the MetaboLights accession numbers REQ20260715221650 and REQ20260715221651 for the NMR and LC-MS data respectively.

### Author Contributions

AGB: Formal analysis, Visualisation, Writing – original draft. KAW: Investigation. GF, JW, EH and JKN: Writing – review and editing. EH and JKN: Funding acquisition

## Supplementary Figures

**Supplementary Figure 1.**
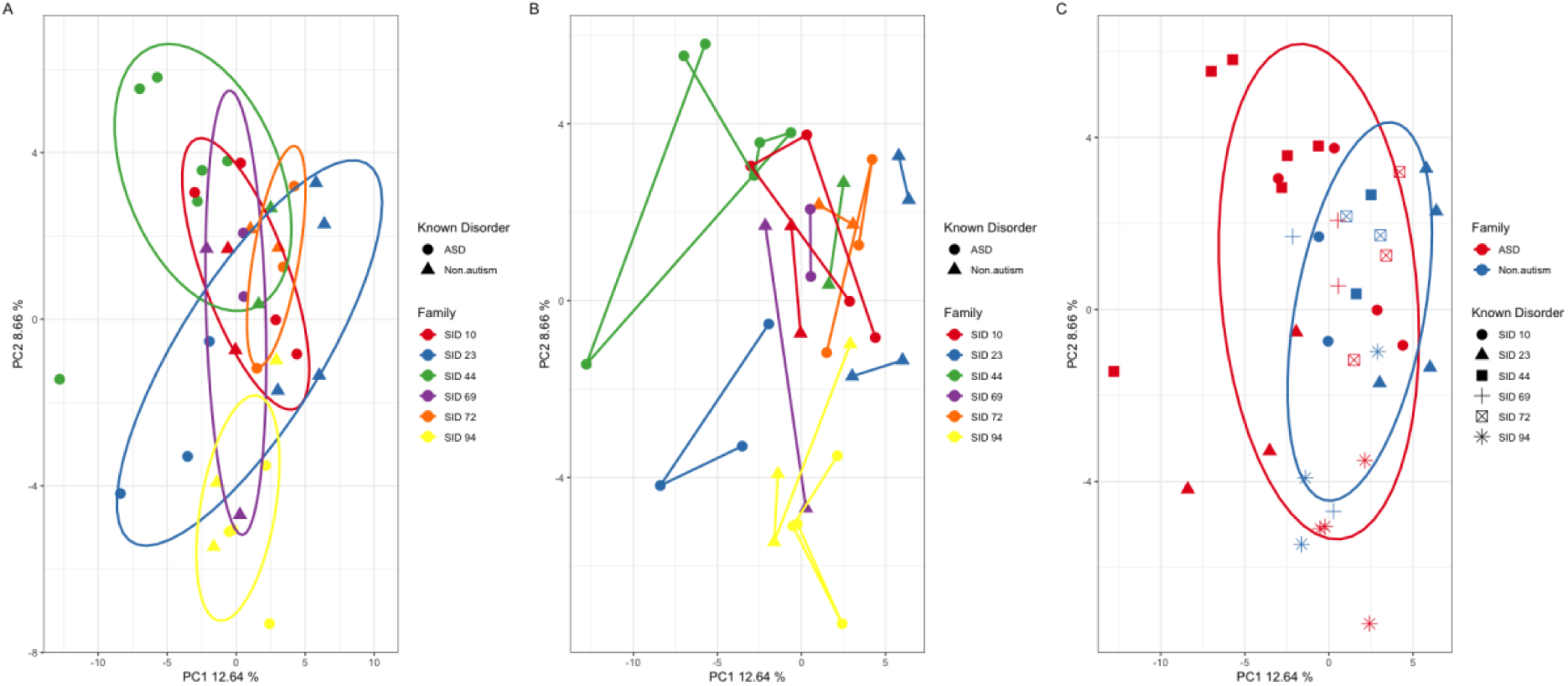
Principal Component Analysis (PCA) of metabolic and microbiome profiles, highlighting the strong influence of family on sample composition. ***(A)*** A PCA plot of the metabolic and microbiome data demonstrates that samples cluster primarily based on their family (indicated by coloured ellipses), with little separation observed between individuals with and without ASD. ***(B)*** Similarly, a PCA of the metabolic and microbiome data shows that individuals’ profile are more similar to themselves over others with the same ASD diagnosis. ***(C)*** When samples are grouped solely by ASD diagnosis (blue) and neurotypical status (red), the PCA reveals a significant overlap between the two ellipses, indicating minimal separation between the groups on the basis of ASD diagnosis alone.

**Supplementary Figure 2.**
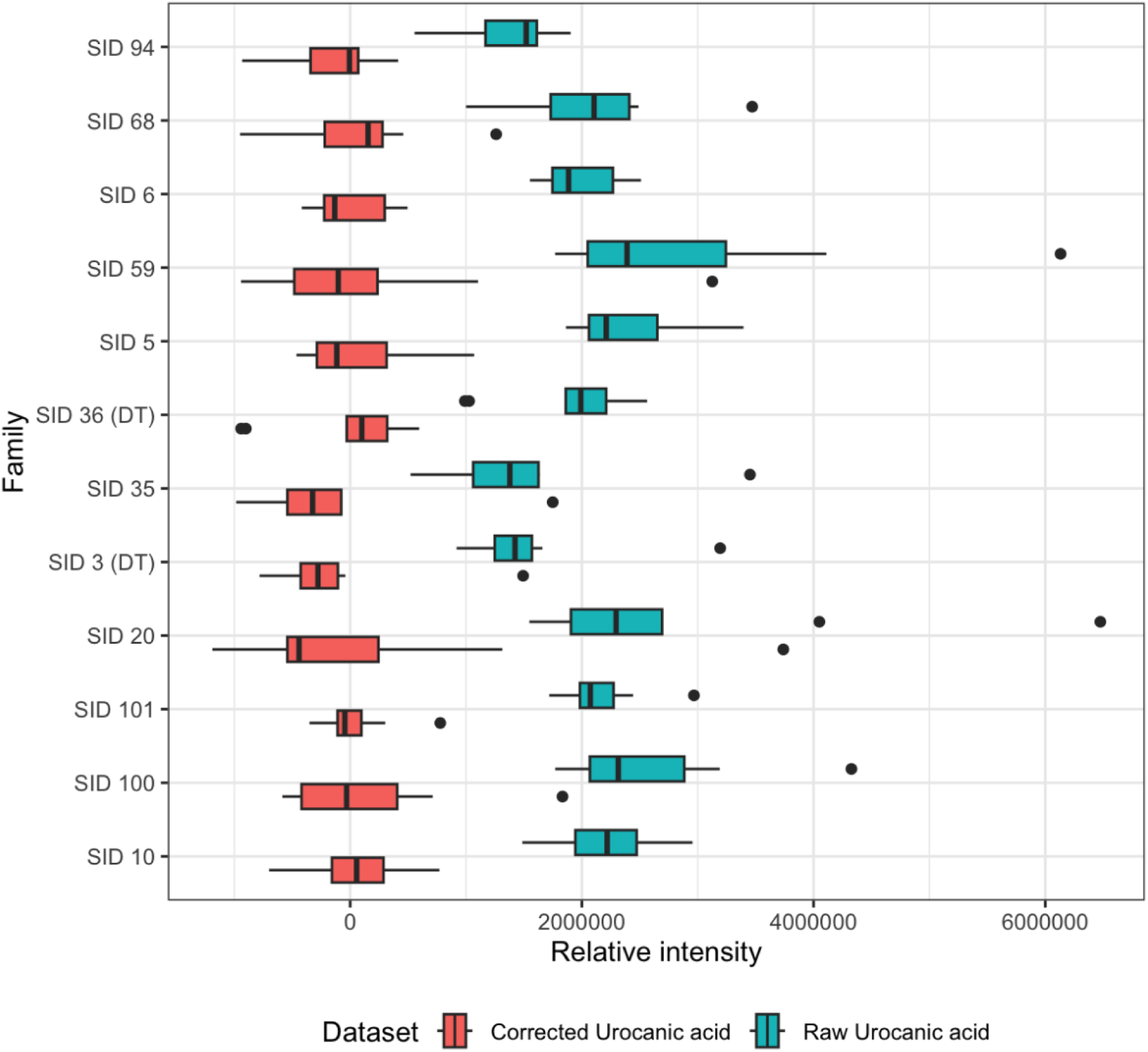
Examples of a family + individual corrected feature (Urocanic acid) (red), versus an uncorrected feature (blue). Formula: resid(Urocanic_acid ∼ (1|family/individual_replicate))

**Supplementary Figure 3.**
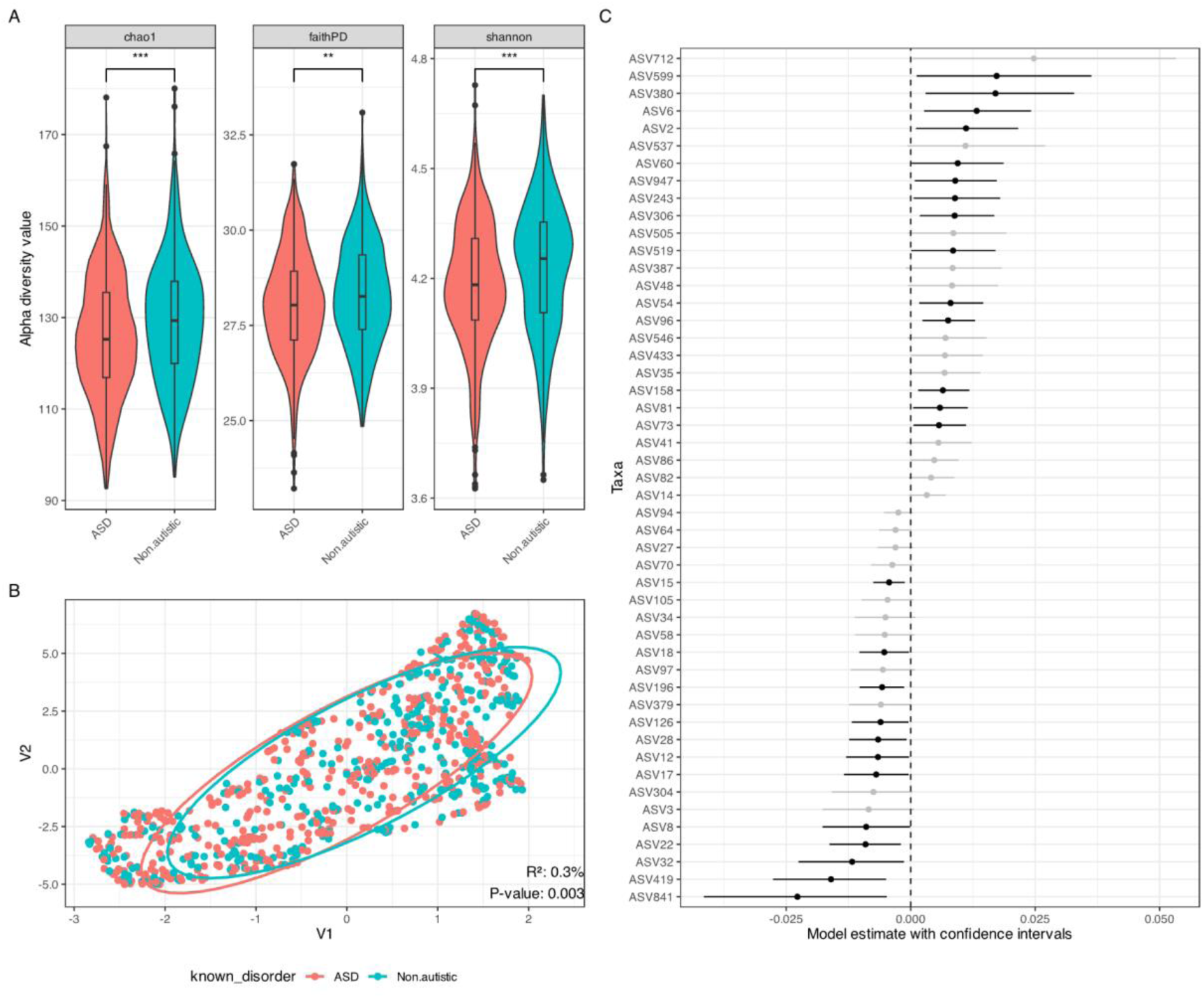
ASV level Microbiome Diversity and Differential Abundance Analysis. *(A)* Alpha diversity (within-sample diversity) of the gut microbiome. Diversity was measured using three metrics: Chao1 (estimating community richness), Faith’s Phylogenetic Diversity (measuring the phylogenetic diversity of taxa), and the Shannon index (quantifying both richness and evenness). *(B)* Beta diversity (between-sample diversity) to show the overall differences in microbial community composition between the groups. The plot is based on Weighted UniFrac, a phylogenetic distance metric that accounts for both the presence and abundance of microbial taxa. *(C)* Linear model used to determine the differential abundance of specific bacterial taxa between the ASD and neurotypical groups. The plot highlights the genera and other taxonomic levels that were found to be statistically significantly altered in the ASD cohort.

**Supplementary Figure 4.**
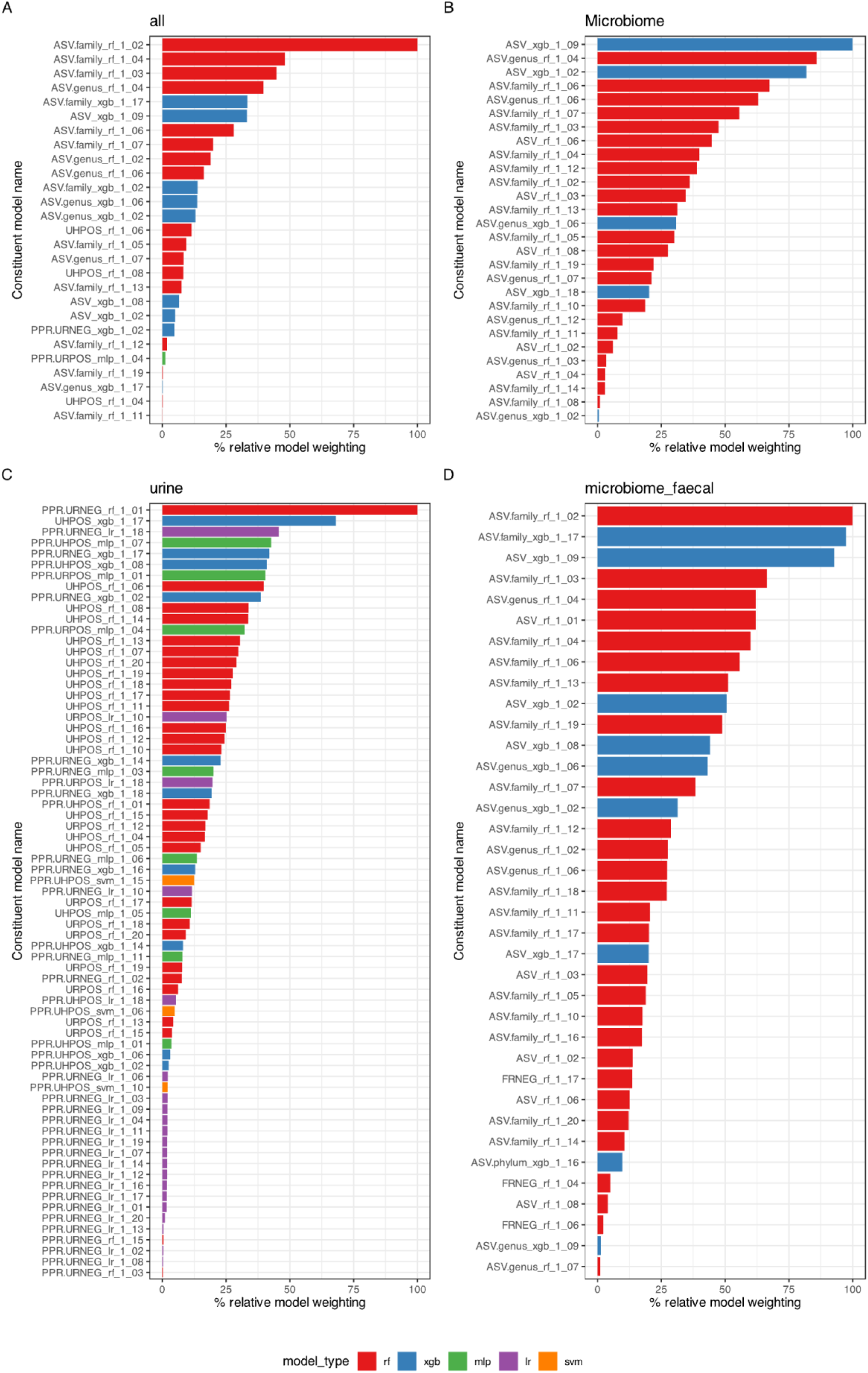
Relative contribution of different classification algorithms to the final ensemble model for four predictive datasets. The percentage contribution of various machine learning models - Random Forest (RF), Extreme Gradient Boosting (XGB), Multi-Layer Perceptron (MLP), Logistic Regression (LR), Support Vector Machine (SVM), k-nearest neighbours (KNN) and naïve bayes (NB) (KNN and NB not shown due to poor predictive performance), and is shown as percentage relative model weighting to the overall predictive power of the ensemble model for each dataset.

**Supplementary Figure 5.**
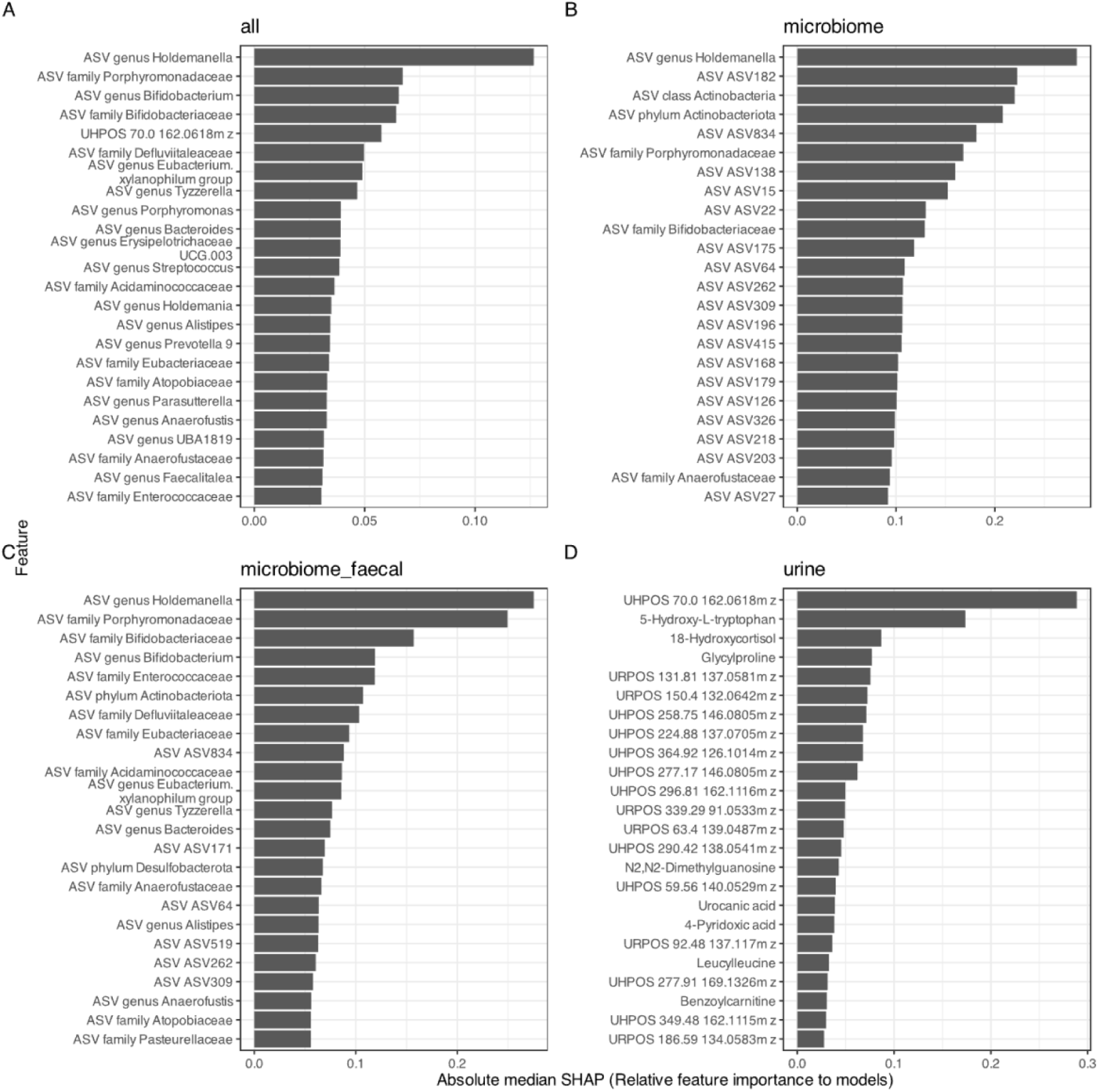
Absolute median SHAP scores of top features with the best ability to classify samples on ASD status

**Supplementary Figure 6.**
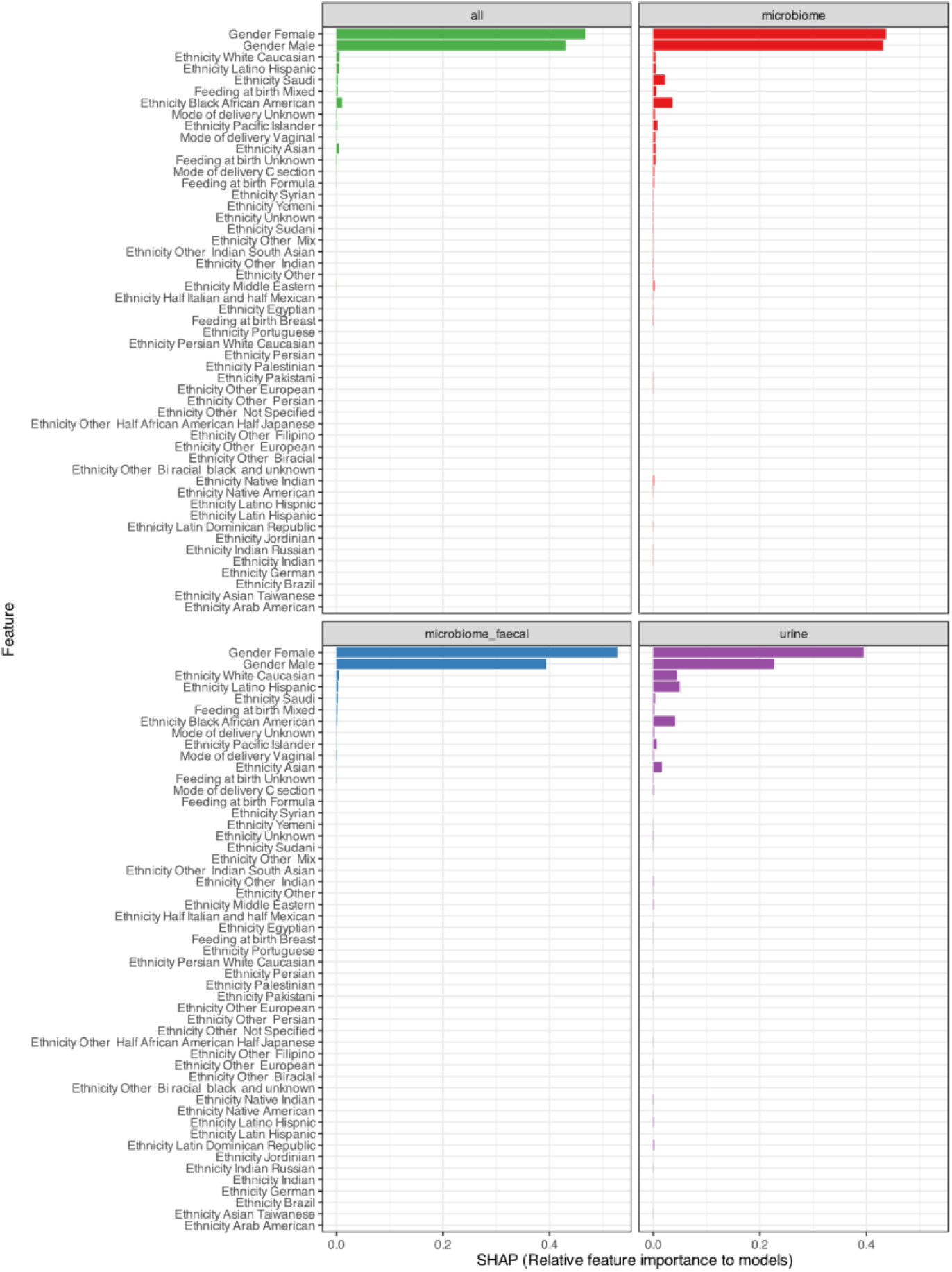
Relative feature importance of biological and lifestyle factors across the ensemble models determined by their contribution to the classification performance of the ensemble models. The importance of these features is shown for different datasets, with Green: The model trained on all datasets, including faecal and urinary metabolites and microbiome taxa. Red: The model trained on microbiome taxa only. Purple: The model trained on urinary metabolites only. Blue: The model trained on microbiome taxa and faecal metabolites.

## Supplementary Tables

**Supplementary Table 1.**
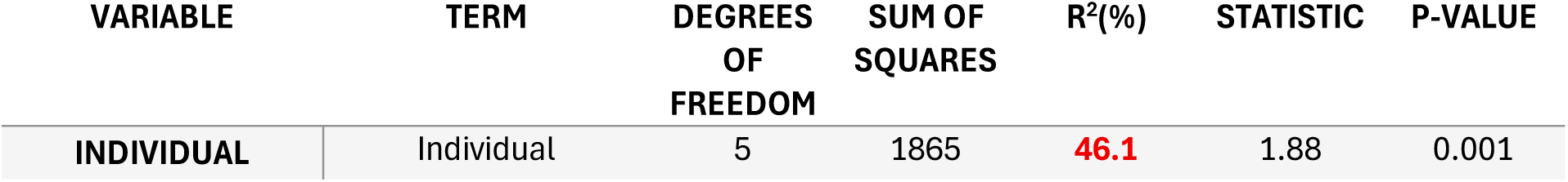
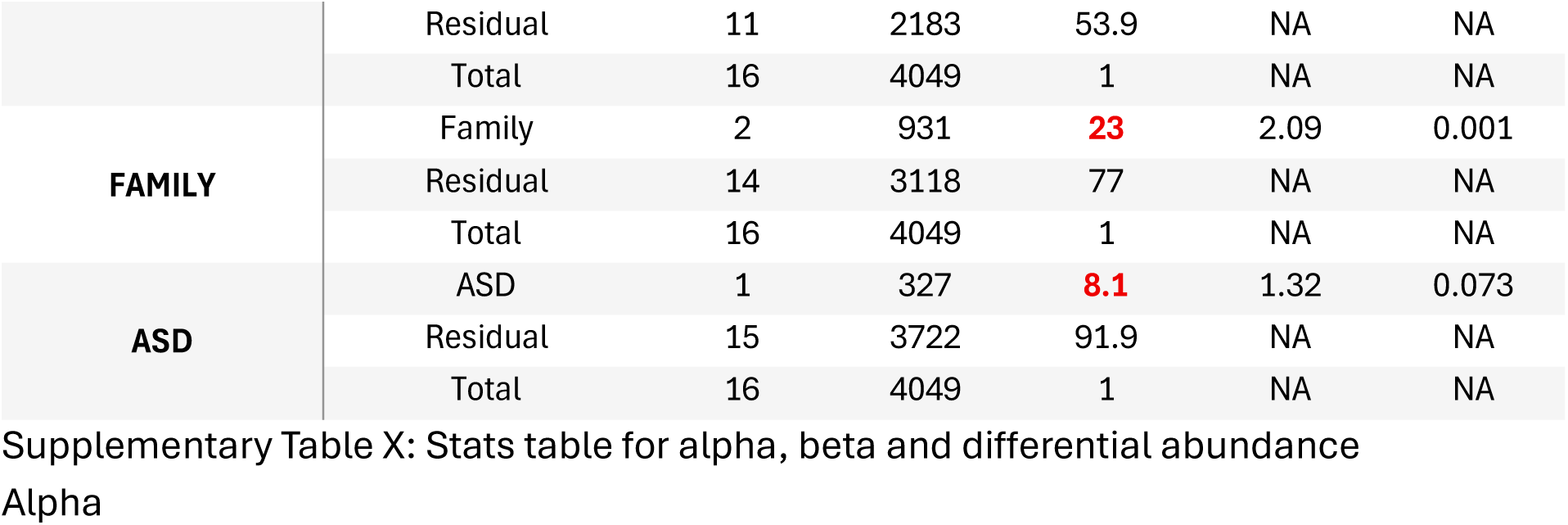
Permutational Multivariate Analysis of Variance (PERMANOVA) quantifying the proportion of total variance explained by different grouping variables within the dataset.

**Supplementary Table 2.**
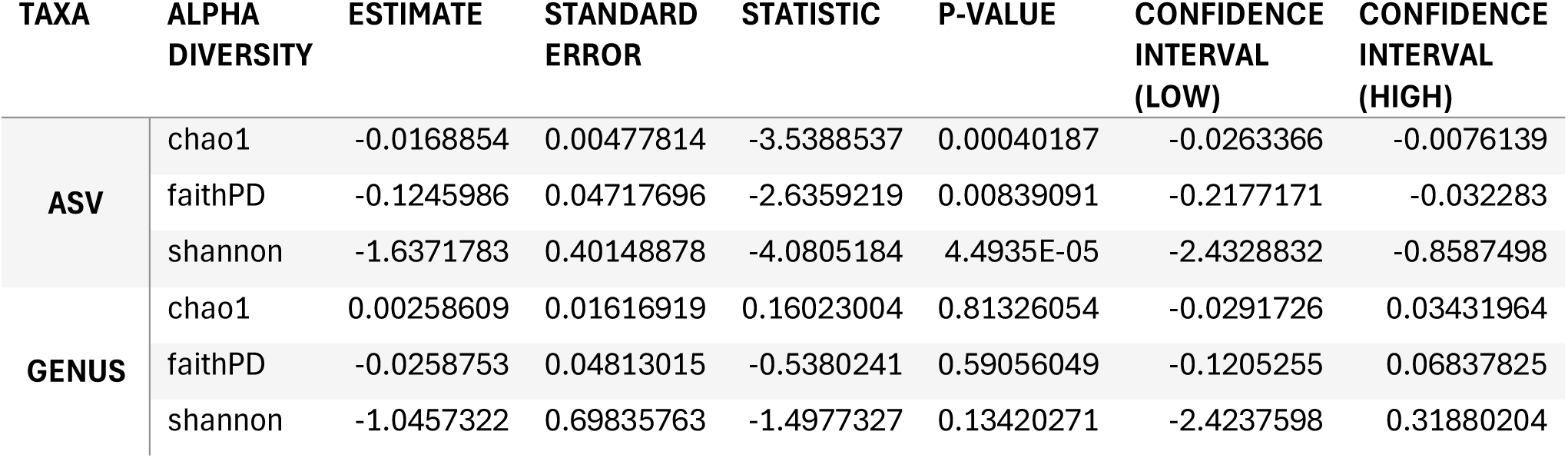
Alpha diversity statistics comparing the diversity of ASD versus neurotypical individuals at an ASD and Genus level

**Supplementary Table 3.**
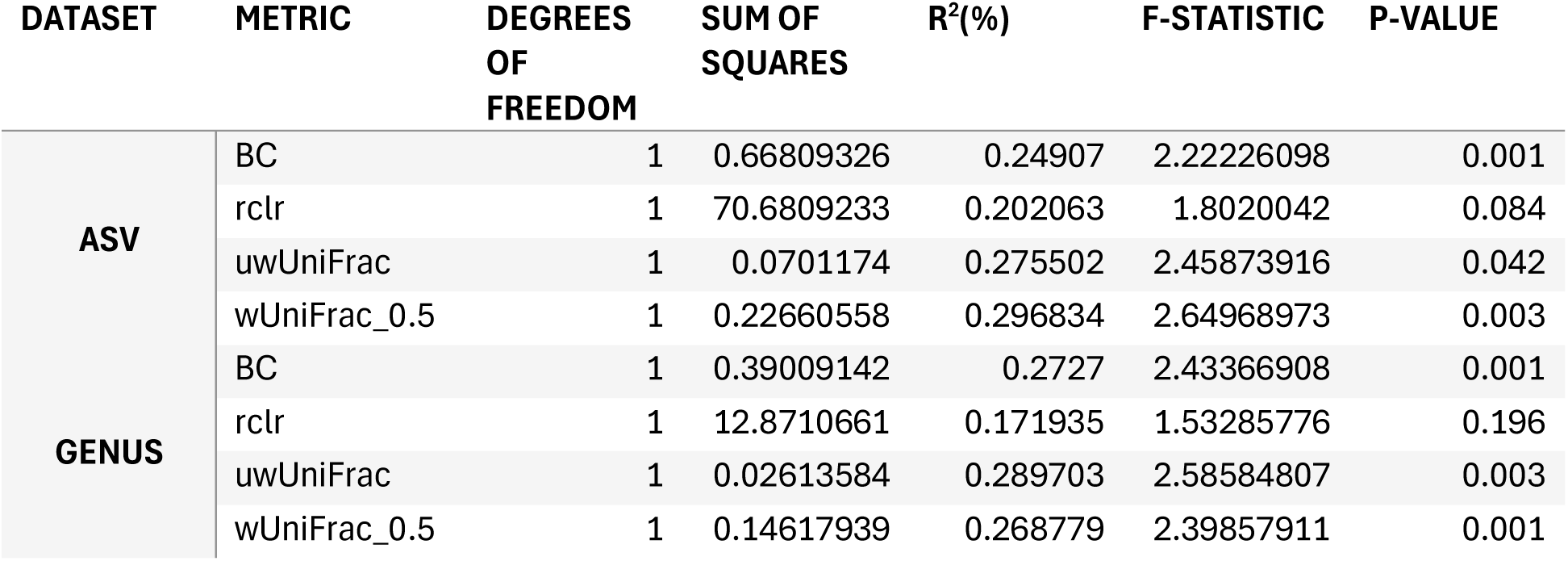
Beta diversity metrics associated with ASD classification as an ASV and Genus taxonomic level

**Supplementary Table 4.**
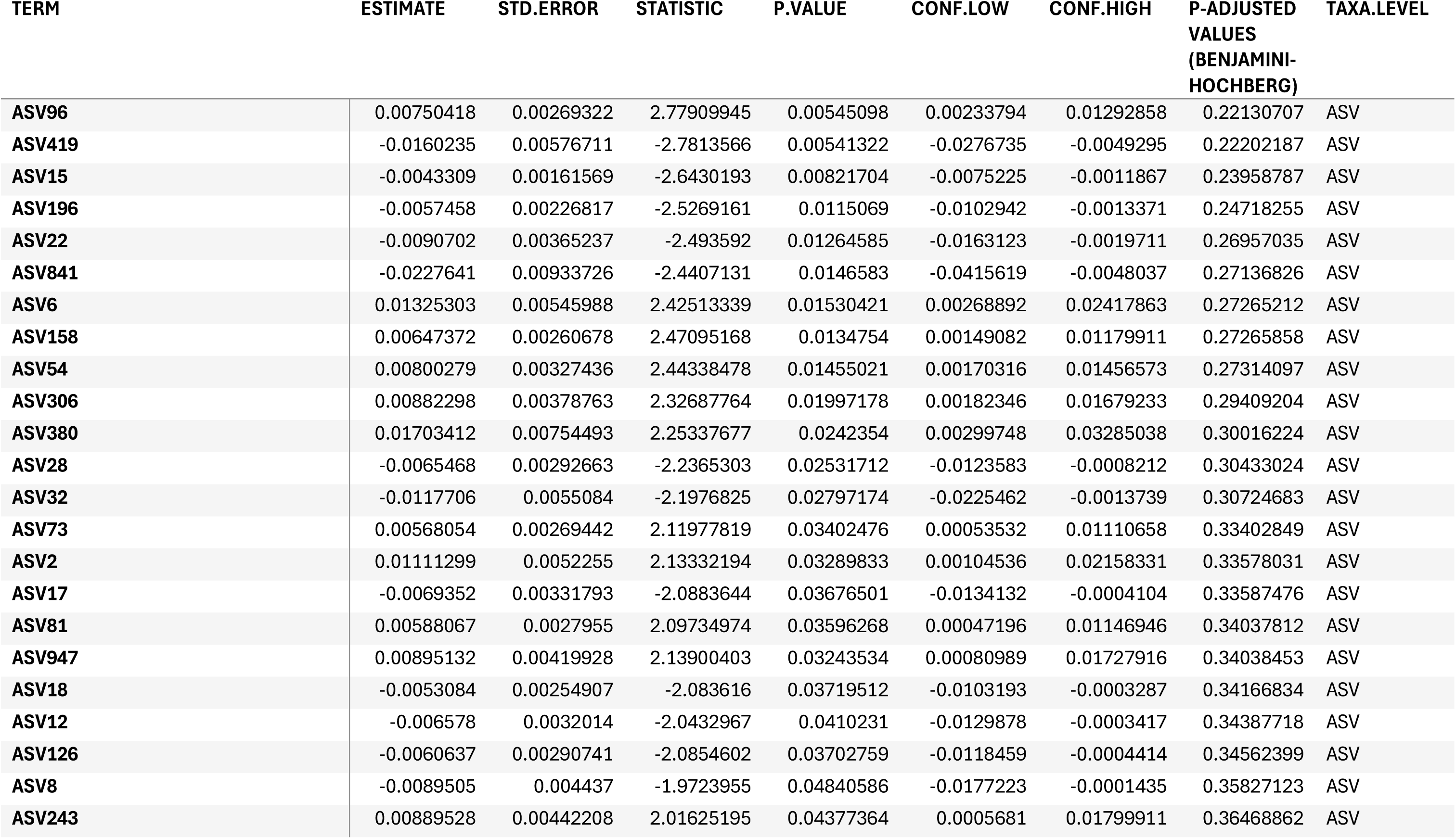
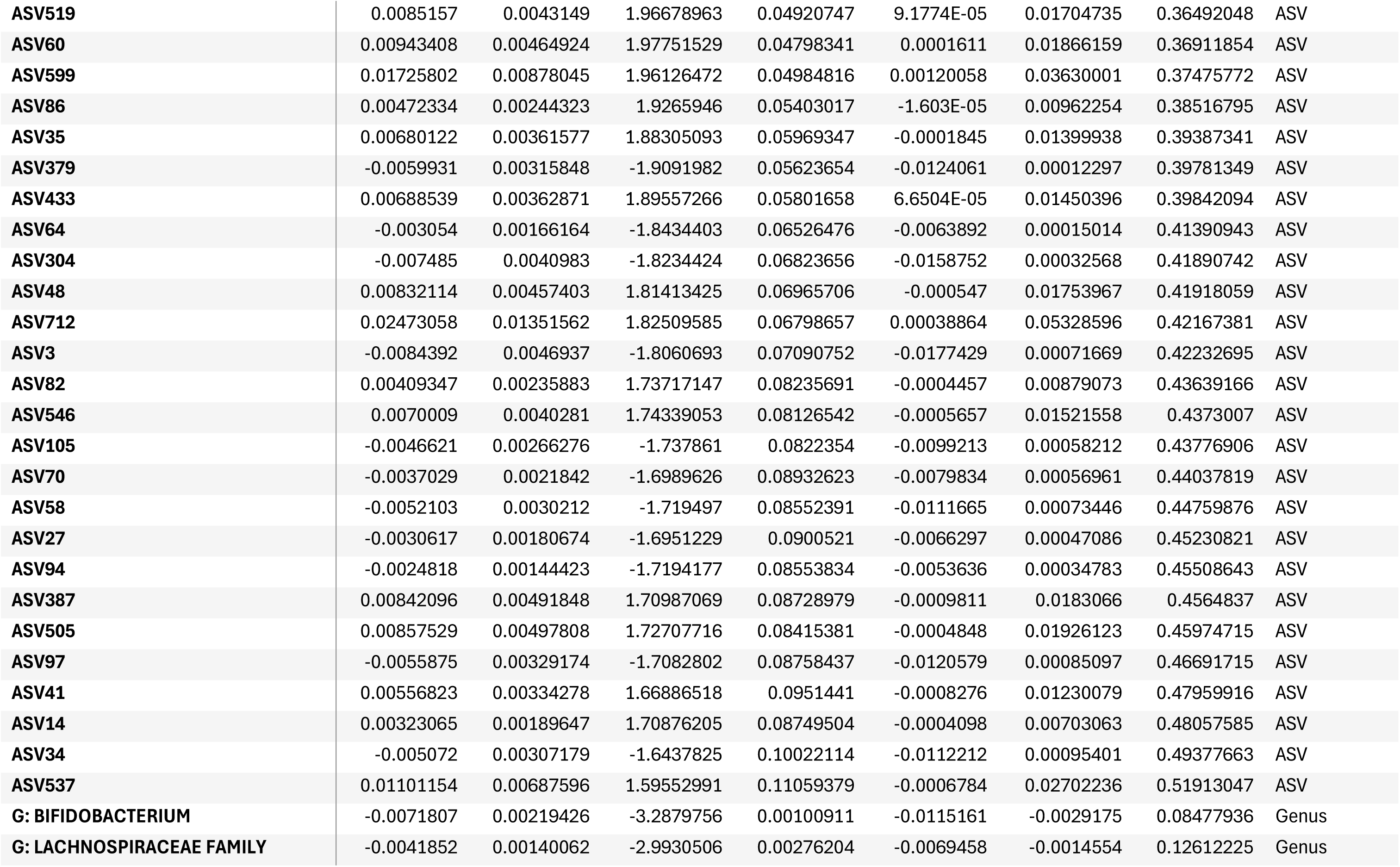
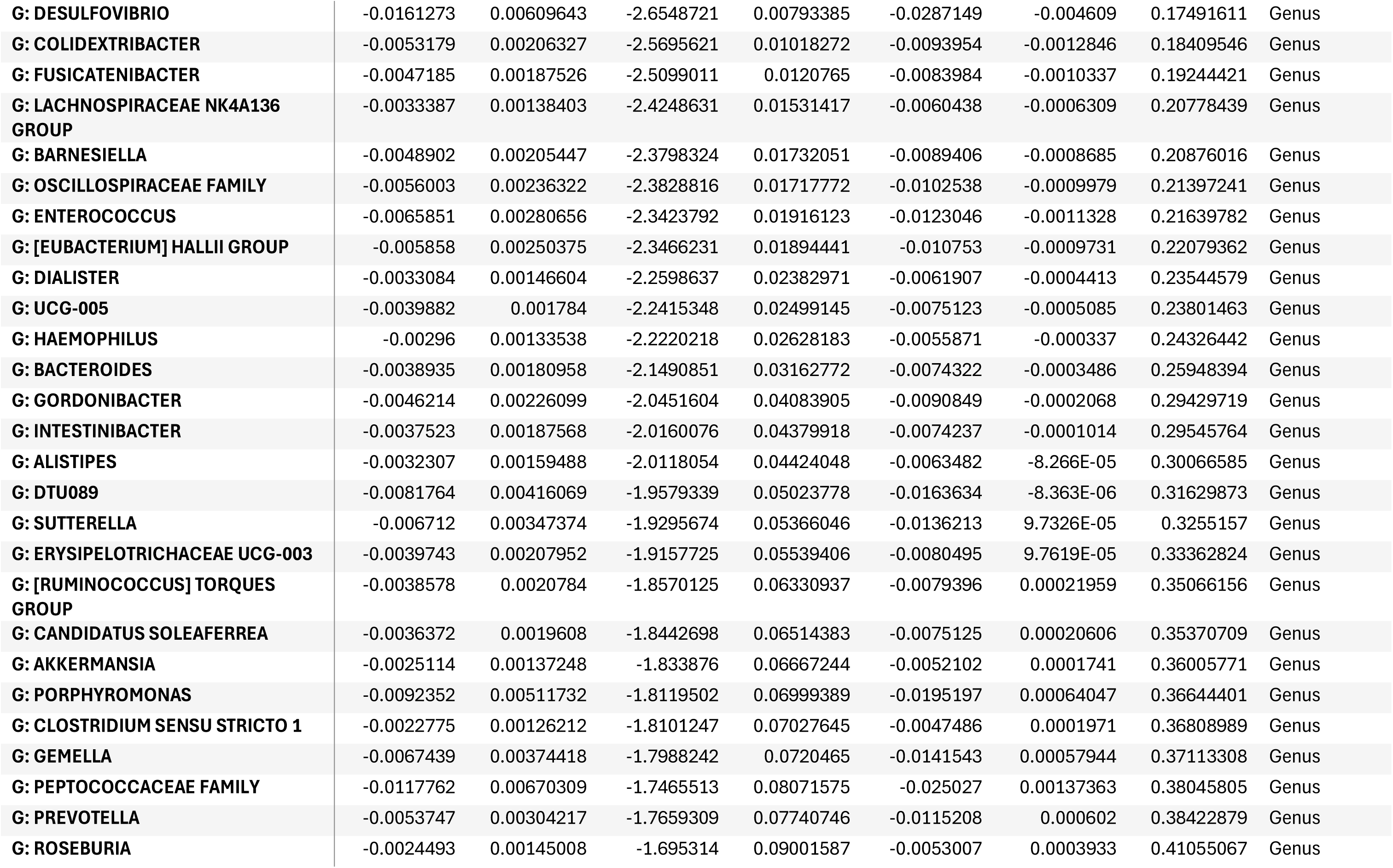
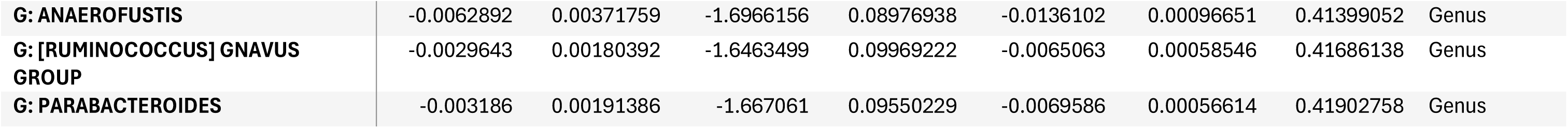
Differential abundance of bacterial taxa associated with ASD status

**Supplementary Table 5.**
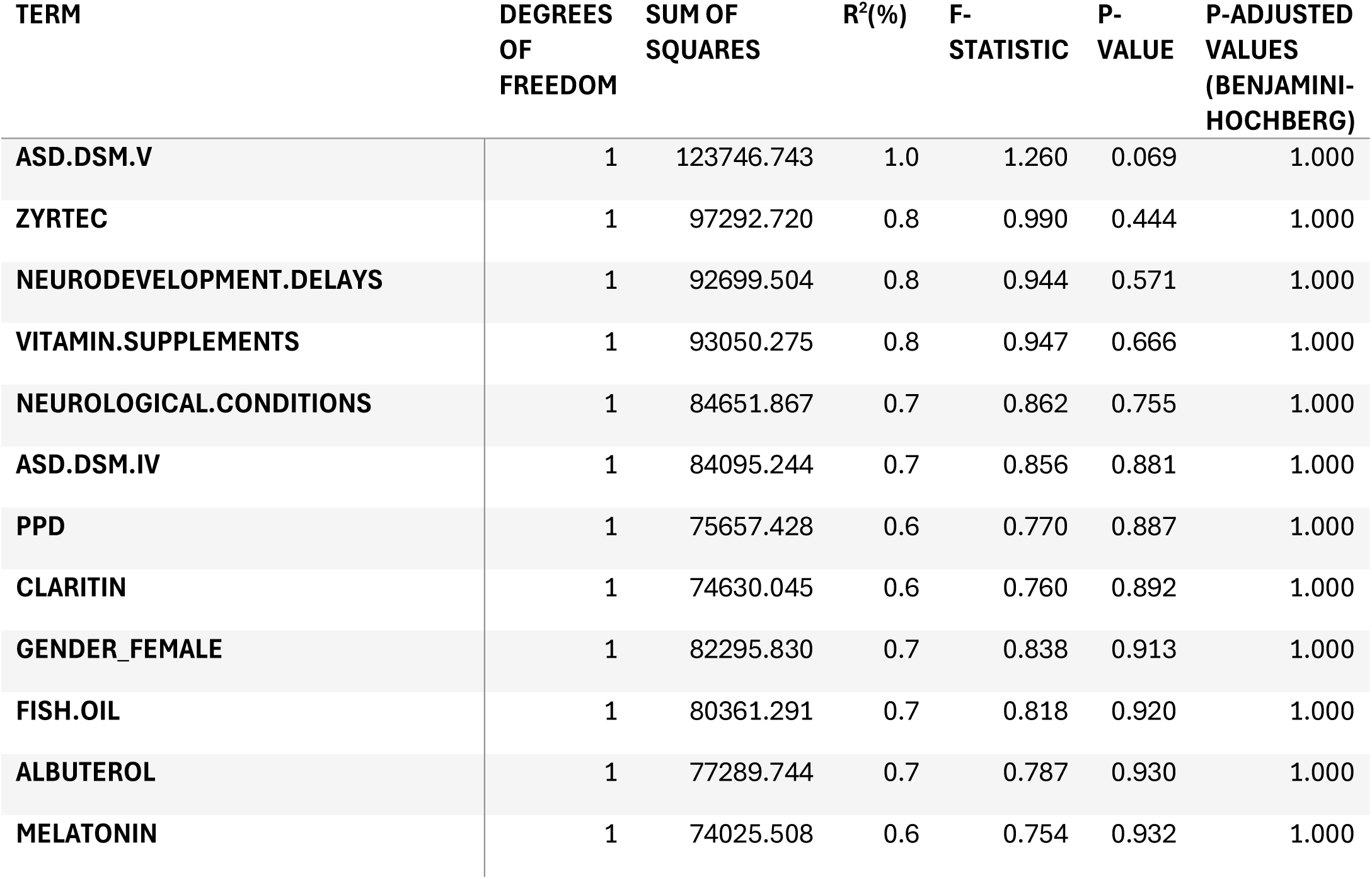
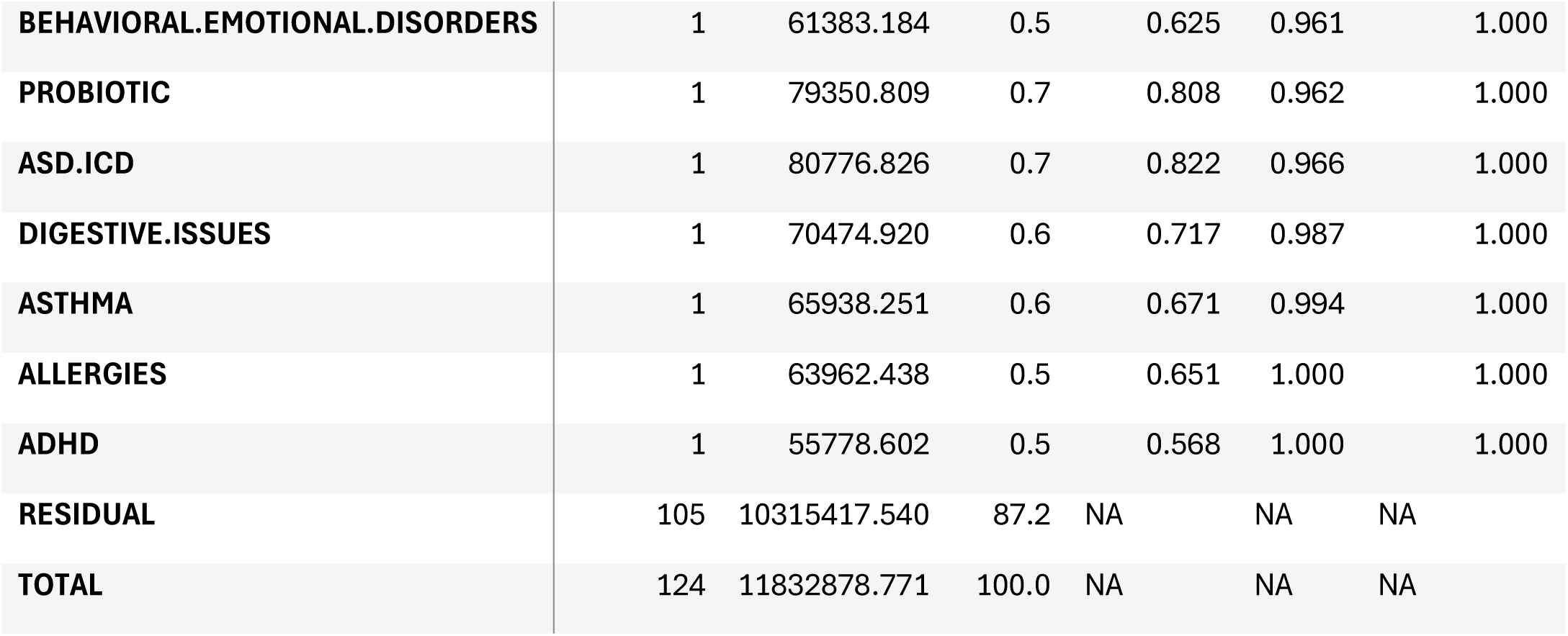
PERMANOVA output of ASD sub-types clustered on Euclidean distances using all datasets combined.

