## Supplementary files for "Multi-Omic Dissection of Autism Reveals Dominant Effects of Family, Sex, and Host-Microbe Metabolic Interactions"

### Supplementary Figures

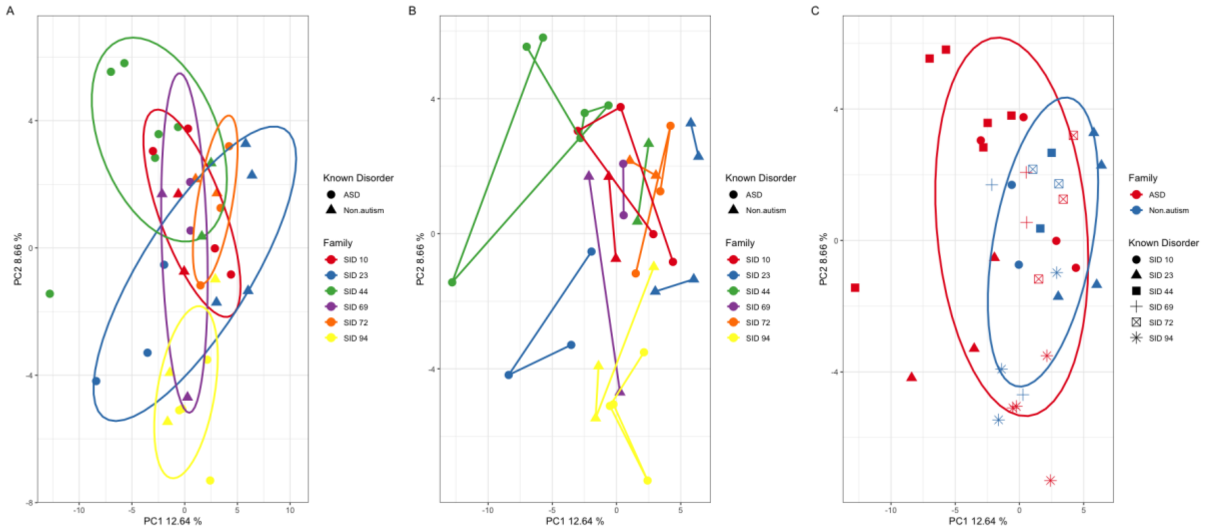


*Supplementary Figure 1 Principal Component Analysis (PCA) of metabolic and microbiome profiles, highlighting the strong influence of family on sample composition.*

***(A)*** *A PCA plot of the metabolic and microbiome data demonstrates that samples cluster primarily based on their family (indicated by coloured ellipses), with little separation observed between individuals with and without ASD.*

***(B)*** *Similarly, a PCA of the metabolic and microbiome data shows that individuals' profile are more similar to themselves over others with the same ASD diagnosis.*

***(C)*** *When samples are grouped solely by ASD diagnosis (blue) and neurotypical status (red), the PCA reveals a significant overlap between the two ellipses, indicating minimal separation between the groups on the basis of ASD diagnosis alone.*

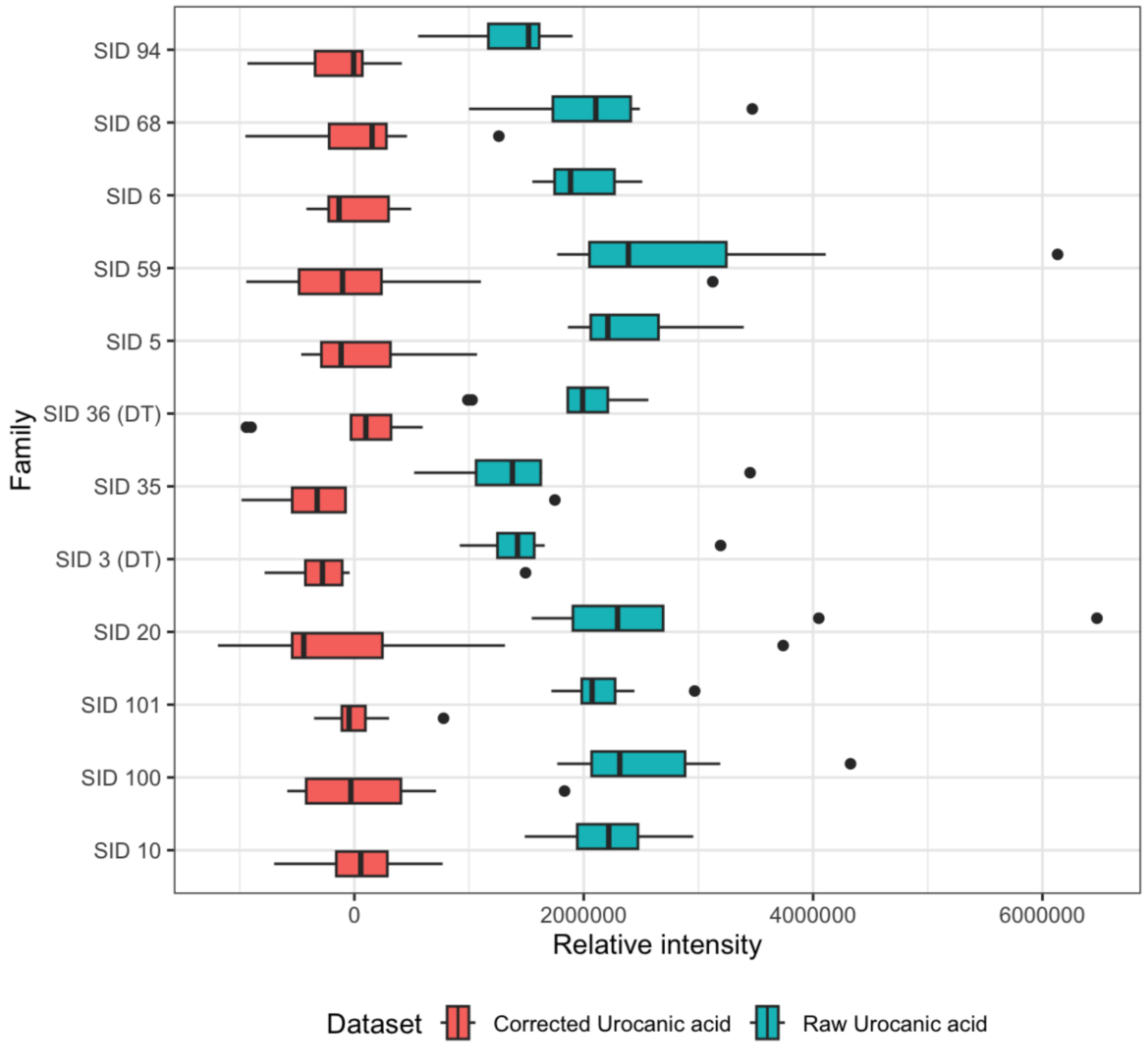


*Supplementary Figure 2 Examples of a family + individual corrected feature (Urocanic acid) (red), versus an uncorrected feature (blue). Formula: resid(Urocanic_acid ~ (1|family/individual_replicate))*

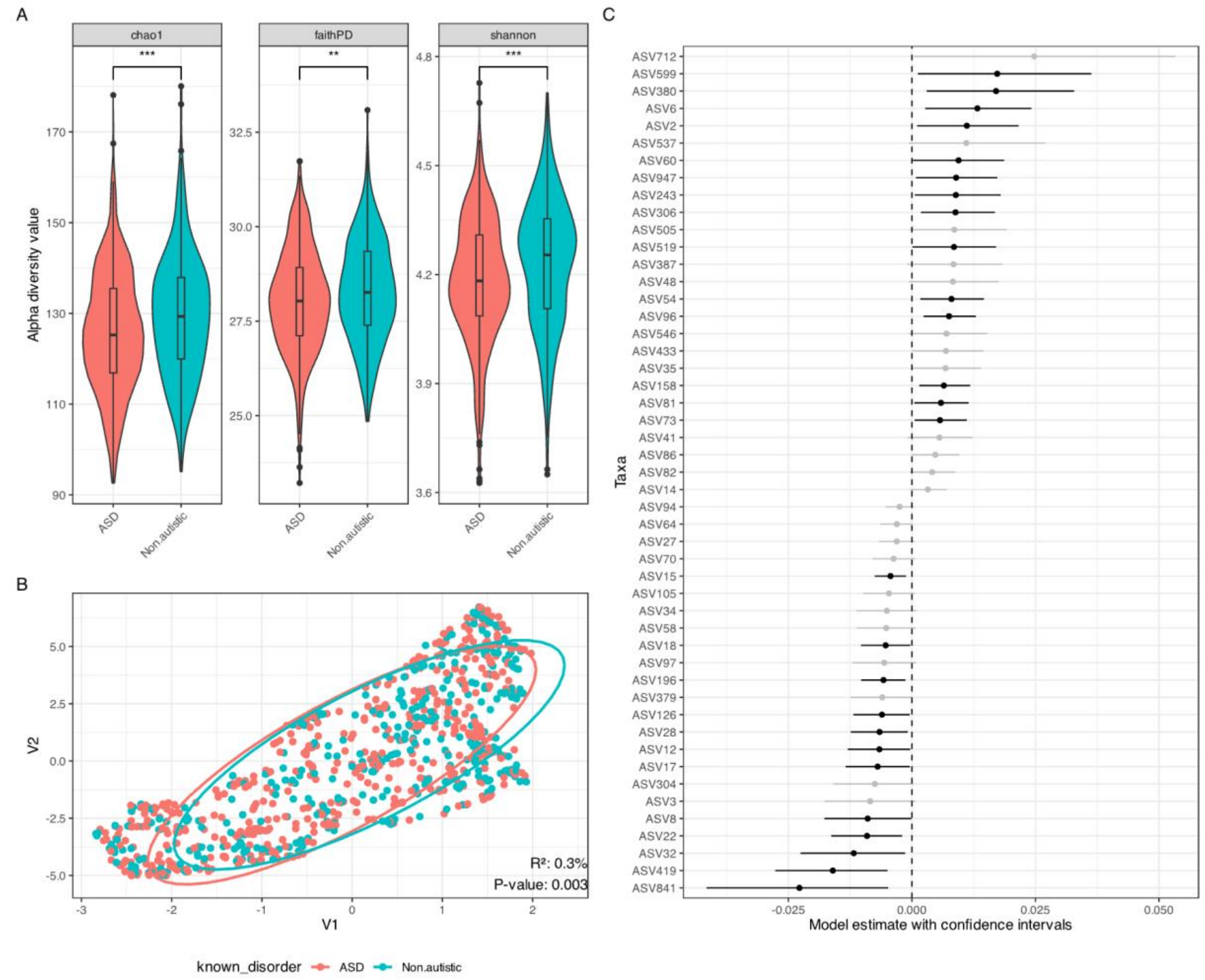


*Supplementary Figure 3 ASV level Microbiome Diversity and Differential Abundance Analysis.*

*(A) Alpha diversity (within-sample diversity) of the gut microbiome. Diversity was measured using three metrics: Chao1 (estimating community richness), Faith's Phylogenetic Diversity (measuring the phylogenetic diversity of taxa), and the Shannon index (quantifying both richness and evenness).*

*(B) Beta diversity (between-sample diversity) to show the overall differences in microbial community composition between the groups. The plot is based on Weighted UniFrac, a phylogenetic distance metric that accounts for both the presence and abundance of microbial taxa.*

*(C) Linear model used to determine the differential abundance of specific bacterial taxa between the ASD and neurotypical groups. The plot highlights the genera and other taxonomic levels that were found to be statistically significantly altered in the ASD cohort.*

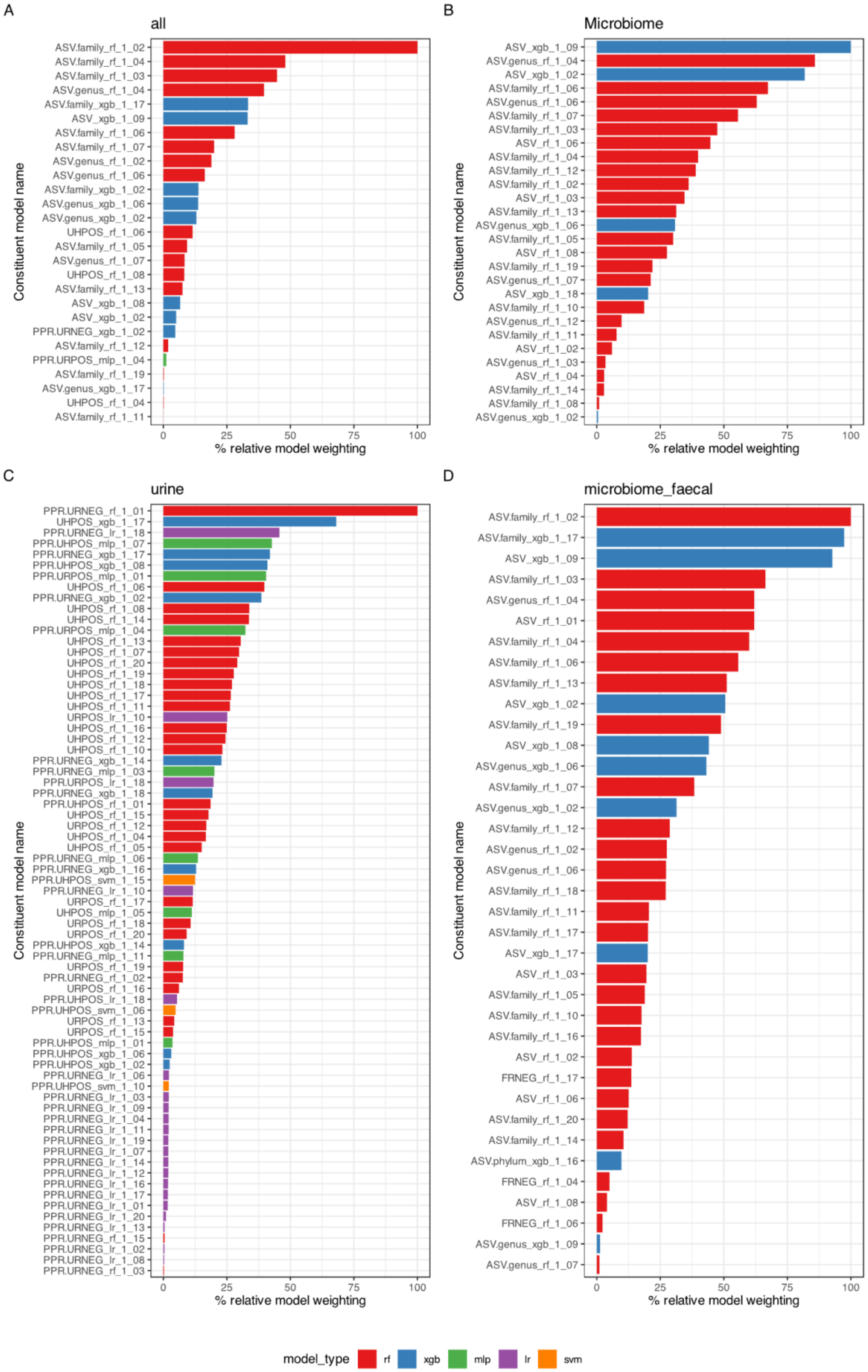
A
all
ASV.family_rf_1_02
ASV.family_rf_1_04
ASV.family_rf_1_03
ASV.genus_rf_1_04
ASV.family_xgb_1_17
ASV_xgb_1_09
ASV.family_rf_1_06
ASV.family_rf_1_07
ASV.genus_rf_1_02
ASV.genus_rf_1_06
ASV.family_xgb_1_02
ASV.genus_xgb_1_06
ASV.genus_xgb_1_02
UHPOS_rf_1_06
ASV.family_rf_1_05
ASV.genus_rf_1_07
UHPOS_rf_1_08
ASV.family_rf_1_13
ASV_xgb_1_08
ASV_xgb_1_02
PPR.URNEG_xgb_1_02
ASV.family_rf_1_12
PPR.URPOS_mlp_1_04
ASV.family_rf_1_19
ASV.genus_xgb_1_17
UHPOS_rf_1_04
ASV.family_rf_1_11
Constituent model name
0
25
50
75
100
% relative model weighting
B
Microbiome
ASV_xgb_1_09
ASV.genus_rf_1_04
ASV_xgb_1_02
ASV.family_rf_1_06
ASV.genus_rf_1_06
ASV.family_rf_1_07
ASV.family_rf_1_03
ASV_rf_1_06
ASV.family_rf_1_04
ASV.family_rf_1_12
ASV.family_rf_1_02
ASV_rf_1_03
ASV.family_rf_1_13
ASV.genus_xgb_1_06
ASV.family_rf_1_05
ASV_rf_1_08
ASV.family_rf_1_19
ASV.genus_rf_1_07
ASV_xgb_1_18
ASV.family_rf_1_10
ASV.genus_rf_1_12
ASV.family_rf_1_11
ASV_rf_1_02
ASV.genus_rf_1_03
ASV_rf_1_04
ASV.family_rf_1_14
ASV.family_rf_1_08
ASV.genus_xgb_1_02
Constituent model name
0
25
50
75
100
% relative model weighting
C
urine
PPR.URNEG_rf_1_01
UHPOS_xgb_1_17
PPR.URNEG_lr_1_18
PPR.UHPOS_mlp_1_07
PPR.URNEG_xgb_1_17
PPR.UHPOS_xgb_1_08
PPR.URPOS_mlp_1_01
UHPOS_rf_1_06
PPR.URNEG_xgb_1_02
UHPOS_rf_1_08
UHPOS_rf_1_14
PPR.URPOS_mlp_1_04
UHPOS_rf_1_13
UHPOS_rf_1_07
UHPOS_rf_1_20
UHPOS_rf_1_19
UHPOS_rf_1_18
UHPOS_rf_1_17
UHPOS_rf_1_11
URPOS_lr_1_10
UHPOS_rf_1_16
UHPOS_rf_1_12
UHPOS_rf_1_10
PPR.URNEG_xgb_1_14
PPR.URNEG_mlp_1_03
PPR.URPOS_lr_1_18
PPR.URNEG_xgb_1_18
PPR.UHPOS_rf_1_01
UHPOS_rf_1_15
URPOS_rf_1_12
UHPOS_rf_1_04
UHPOS_rf_1_05
PPR.URNEG_mlp_1_06
PPR.URNEG_xgb_1_16
PPR.UHPOS_svm_1_15
PPR.URNEG_lr_1_10
URPOS_rf_1_17
UHPOS_mlp_1_05
URPOS_rf_1_18
URPOS_rf_1_20
PPR.UHPOS_xgb_1_14
PPR.URNEG_mlp_1_11
URPOS_rf_1_19
PPR.URNEG_rf_1_02
URPOS_rf_1_16
PPR.UHPOS_lr_1_18
PPR.UHPOS_svm_1_06
URPOS_rf_1_13
URPOS_rf_1_15
PPR.UHPOS_mlp_1_01
PPR.UHPOS_xgb_1_06
PPR.UHPOS_xgb_1_02
PPR.URNEG_lr_1_06
PPR.UHPOS_svm_1_10
PPR.URNEG_lr_1_03
PPR.URNEG_lr_1_09
PPR.URNEG_lr_1_04
PPR.URNEG_lr_1_11
PPR.URNEG_lr_1_19
PPR.URNEG_lr_1_07
PPR.URNEG_lr_1_14
PPR.URNEG_lr_1_12
PPR.URNEG_lr_1_16
PPR.URNEG_lr_1_17
PPR.URNEG_lr_1_01
PPR.URNEG_lr_1_20
PPR.URNEG_lr_1_13
PPR.URNEG_rf_1_15
PPR.URNEG_lr_1_02
PPR.URNEG_lr_1_08
PPR.URNEG_rf_1_03
Constituent model name
0
25
50
75
100
% relative model weighting
D
microbiome_faecal
ASV.family_rf_1_02
ASV.family_xgb_1_17
ASV_xgb_1_09
ASV.family_rf_1_03
ASV.genus_rf_1_04
ASV_rf_1_01
ASV.family_rf_1_04
ASV.family_rf_1_06
ASV.family_rf_1_13
ASV_xgb_1_02
ASV.family_rf_1_19
ASV_xgb_1_08
ASV.genus_xgb_1_06
ASV.family_rf_1_07
ASV.genus_xgb_1_02
ASV.family_rf_1_12
ASV.genus_rf_1_02
ASV.genus_rf_1_06
ASV.family_rf_1_18
ASV.family_rf_1_11
ASV.family_rf_1_17
ASV_xgb_1_17
ASV_rf_1_03
ASV.family_rf_1_05
ASV.family_rf_1_10
ASV.family_rf_1_16
ASV_rf_1_02
FRNEG_rf_1_17
ASV_rf_1_06
ASV.family_rf_1_20
ASV.family_rf_1_14
ASV.phylum_xgb_1_16
FRNEG_rf_1_04
ASV_rf_1_08
FRNEG_rf_1_06
ASV.genus_xgb_1_09
ASV.genus_rf_1_07
Constituent model name
0
25
50
75
100
% relative model weighting
model_type
rf
xgb
mlp
lr
svm

*Supplementary Figure 4 Relative contribution of different classification algorithms to the final ensemble model for four predictive datasets. The percentage contribution of various machine learning models - Random Forest (RF), Extreme Gradient Boosting (XGB), Multi-Layer Perceptron (MLP), Logistic Regression (LR), Support Vector Machine (SVM), k-nearest neighbours (KNN) and naïve bayes (NB)  (KNN and NB not shown due to poor predictive performance), and is shown as percentage relative model weighting to the overall predictive power of the ensemble model for each dataset.*

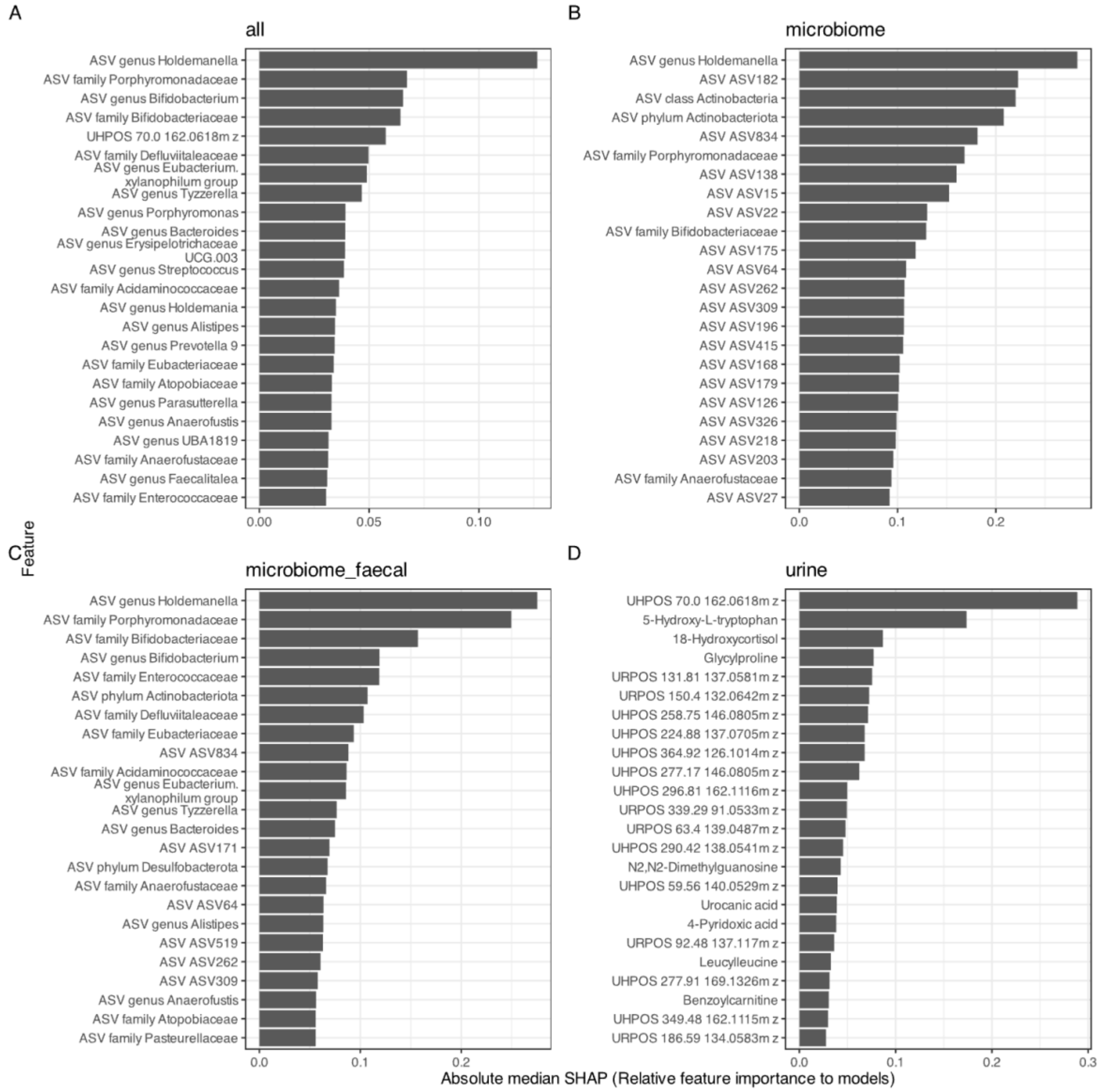


*Supplementary Figure 5 Absolute median SHAP scores of top features with the best ability to classify samples on ASD status*

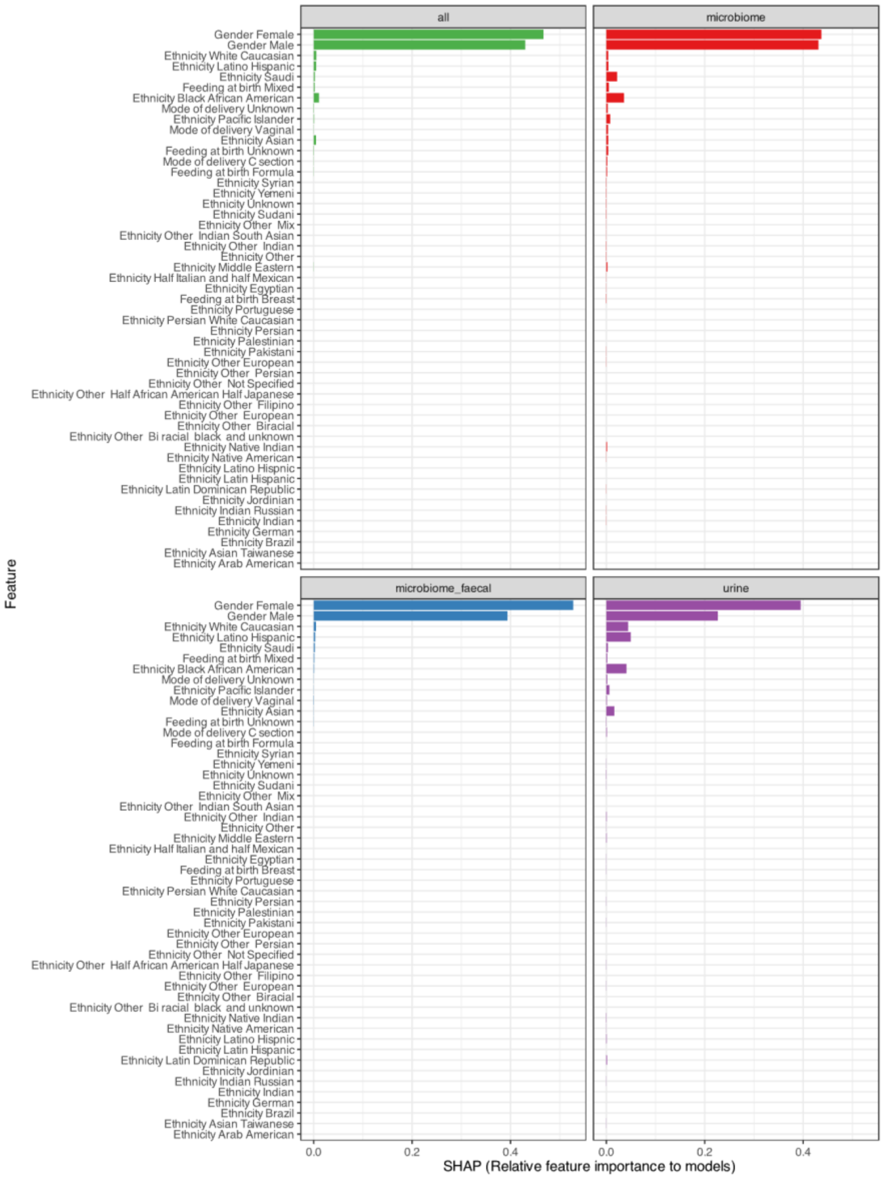


*Supplementary Figure 6 Relative feature importance of biological and lifestyle factors across the ensemble models determined by their contribution to the classification performance of the ensemble models. The importance of these features is shown for different datasets, with Green: The model trained on all datasets, including faecal and urinary metabolites and microbiome taxa. Red: The model trained on microbiome taxa only. Purple: The model trained on urinary metabolites only. Blue: The model trained on microbiome taxa and faecal metabolites.*

### Supplementary Tables

*Supplementary Table 1 Permutational Multivariate Analysis of Variance (PERMANOVA) quantifying the proportion of total variance explained by different grouping variables within the dataset.*

| VARIABLE | TERM | DEGREES OF FREEDOM | SUM OF SQUARES | $R^2$(%) | STATISTIC | P-VALUE |
|---|---|---|---|---|---|---|
| **INDIVIDUAL** | Individual | 5 | 1865 | **46.1** | 1.88 | 0.001 |

| | | | | | | |
|---|---|---|---|---|---|---|
| | Residual | 11 | 2183 | 53.9 | NA | NA |
| | Total | 16 | 4049 | 1 | NA | NA |
| **FAMILY** | Family | 2 | 931 | **23** | 2.09 | 0.001 |
| | Residual | 14 | 3118 | 77 | NA | NA |
| | Total | 16 | 4049 | 1 | NA | NA |
| **ASD** | ASD | 1 | 327 | **8.1** | 1.32 | 0.073 |
| | Residual | 15 | 3722 | 91.9 | NA | NA |
| | Total | 16 | 4049 | 1 | NA | NA |

Supplementary Table X: Stats table for alpha, beta and differential abundance

Alpha

*Supplementary Table 2 Alpha diversity statistics comparing the diversity of ASD versus neurotypical individuals at an ASD and Genus level*

| TAXA | ALPHA DIVERSITY | ESTIMATE | STANDARD ERROR | STATISTIC | P-VALUE | CONFIDENCE INTERVAL (LOW) | CONFIDENCE INTERVAL (HIGH) |
|---|---|---|---|---|---|---|---|
| **ASV** | chao1 | -0.0168854 | 0.00477814 | -3.5388537 | 0.00040187 | -0.0263366 | -0.0076139 |
| | faithPD | -0.1245986 | 0.04717696 | -2.6359219 | 0.00839091 | -0.2177171 | -0.032283 |
| | shannon | -1.6371783 | 0.40148878 | -4.0805184 | 4.4935E-05 | -2.4328832 | -0.8587498 |
| **GENUS** | chao1 | 0.00258609 | 0.01616919 | 0.16023004 | 0.81326054 | -0.0291726 | 0.03431964 |
| | faithPD | -0.0258753 | 0.04813015 | -0.5380241 | 0.59056049 | -0.1205255 | 0.06837825 |
| | shannon | -1.0457322 | 0.69835763 | -1.4977327 | 0.13420271 | -2.4237598 | 0.31880204 |

*Supplementary Table 3 Beta diversity metrics associated with ASD classification as an ASV and Genus taxonomic level*

| DATASET | METRIC | DEGREES OF FREEDOM | SUM OF SQUARES | $R^2$(%) | F-STATISTIC | P-VALUE |
|---|---|---|---|---|---|---|
| **ASV** | BC | 1 | 0.66809326 | 0.24907 | 2.22226098 | 0.001 |
| | rclr | 1 | 70.6809233 | 0.202063 | 1.8020042 | 0.084 |
| | uwUniFrac | 1 | 0.0701174 | 0.275502 | 2.45873916 | 0.042 |
| | wUniFrac_0.5 | 1 | 0.22660558 | 0.296834 | 2.64968973 | 0.003 |
| **GENUS** | BC | 1 | 0.39009142 | 0.2727 | 2.43366908 | 0.001 |
| | rclr | 1 | 12.8710661 | 0.171935 | 1.53285776 | 0.196 |
| | uwUniFrac | 1 | 0.02613584 | 0.289703 | 2.58584807 | 0.003 |
| | wUniFrac_0.5 | 1 | 0.14617939 | 0.268779 | 2.39857911 | 0.001 |

*Supplementary Table 4 Differential abundance of bacterial taxa associated with ASD status*

| TERM | ESTIMATE | STD.ERROR | STATISTIC | P.VALUE | CONF.LOW | CONF.HIGH | P-ADJUSTED VALUES (BENJAMINI-HOCHBERG) | TAXA.LEVEL |
|---|---|---|---|---|---|---|---|---|
| **ASV96** | 0.00750418 | 0.00269322 | 2.77909945 | 0.00545098 | 0.00233794 | 0.01292858 | 0.22130707 | ASV |
| **ASV419** | -0.0160235 | 0.00576711 | -2.7813566 | 0.00541322 | -0.0276735 | -0.0049295 | 0.22202187 | ASV |
| **ASV15** | -0.0043309 | 0.00161569 | -2.6430193 | 0.00821704 | -0.0075225 | -0.0011867 | 0.23958787 | ASV |
| **ASV196** | -0.0057458 | 0.00226817 | -2.5269161 | 0.0115069 | -0.0102942 | -0.0013371 | 0.24718255 | ASV |
| **ASV22** | -0.0090702 | 0.00365237 | -2.493592 | 0.01264585 | -0.0163123 | -0.0019711 | 0.26957035 | ASV |
| **ASV841** | -0.0227641 | 0.00933726 | -2.4407131 | 0.0146583 | -0.0415619 | -0.0048037 | 0.27136826 | ASV |
| **ASV6** | 0.01325303 | 0.00545988 | 2.42513339 | 0.01530421 | 0.00268892 | 0.02417863 | 0.27265212 | ASV |
| **ASV158** | 0.00647372 | 0.00260678 | 2.47095168 | 0.0134754 | 0.00149082 | 0.01179911 | 0.27265858 | ASV |
| **ASV54** | 0.00800279 | 0.00327436 | 2.44338478 | 0.01455021 | 0.00170316 | 0.01456573 | 0.27314097 | ASV |
| **ASV306** | 0.00882298 | 0.00378763 | 2.32687764 | 0.01997178 | 0.00182346 | 0.01679233 | 0.29409204 | ASV |
| **ASV380** | 0.01703412 | 0.00754493 | 2.25337677 | 0.0242354 | 0.00299748 | 0.03285038 | 0.30016224 | ASV |
| **ASV28** | -0.0065468 | 0.00292663 | -2.2365303 | 0.02531712 | -0.0123583 | -0.0008212 | 0.30433024 | ASV |
| **ASV32** | -0.0117706 | 0.0055084 | -2.1976825 | 0.02797174 | -0.0225462 | -0.0013739 | 0.30724683 | ASV |
| **ASV73** | 0.00568054 | 0.00269442 | 2.11977819 | 0.03402476 | 0.00053532 | 0.01110658 | 0.33402849 | ASV |
| **ASV2** | 0.01111299 | 0.0052255 | 2.13332194 | 0.03289833 | 0.00104536 | 0.02158331 | 0.33578031 | ASV |
| **ASV17** | -0.0069352 | 0.00331793 | -2.0883644 | 0.03676501 | -0.0134132 | -0.0004104 | 0.33587476 | ASV |
| **ASV81** | 0.00588067 | 0.0027955 | 2.09734974 | 0.03596268 | 0.00047196 | 0.01146946 | 0.34037812 | ASV |
| **ASV947** | 0.00895132 | 0.00419928 | 2.13900403 | 0.03243534 | 0.00080989 | 0.01727916 | 0.34038453 | ASV |
| **ASV18** | -0.0053084 | 0.00254907 | -2.083616 | 0.03719512 | -0.0103193 | -0.0003287 | 0.34166834 | ASV |
| **ASV12** | -0.006578 | 0.0032014 | -2.0432967 | 0.0410231 | -0.0129878 | -0.0003417 | 0.34387718 | ASV |
| **ASV126** | -0.0060637 | 0.00290741 | -2.0854602 | 0.03702759 | -0.0118459 | -0.0004414 | 0.34562399 | ASV |
| **ASV8** | -0.0089505 | 0.004437 | -1.9723955 | 0.04840586 | -0.0177223 | -0.0001435 | 0.35827123 | ASV |
| **ASV243** | 0.00889528 | 0.00442208 | 2.01625195 | 0.04377364 | 0.0005681 | 0.01799911 | 0.36468862 | ASV |

| | | | | | | | | |
|---|---|---|---|---|---|---|---|---|
| **ASV519** | 0.0085157 | 0.0043149 | 1.96678963 | 0.04920747 | 9.1774E-05 | 0.01704735 | 0.36492048 | ASV |
| **ASV60** | 0.00943408 | 0.00464924 | 1.97751529 | 0.04798341 | 0.0001611 | 0.01866159 | 0.36911854 | ASV |
| **ASV599** | 0.01725802 | 0.00878045 | 1.96126472 | 0.04984816 | 0.00120058 | 0.03630001 | 0.37475772 | ASV |
| **ASV86** | 0.00472334 | 0.00244323 | 1.9265946 | 0.05403017 | -1.603E-05 | 0.00962254 | 0.38516795 | ASV |
| **ASV35** | 0.00680122 | 0.00361577 | 1.88305093 | 0.05969347 | -0.0001845 | 0.01399938 | 0.39387341 | ASV |
| **ASV379** | -0.0059931 | 0.00315848 | -1.9091982 | 0.05623654 | -0.0124061 | 0.00012297 | 0.39781349 | ASV |
| **ASV433** | 0.00688539 | 0.00362871 | 1.89557266 | 0.05801658 | 6.6504E-05 | 0.01450396 | 0.39842094 | ASV |
| **ASV64** | -0.003054 | 0.00166164 | -1.8434403 | 0.06526476 | -0.0063892 | 0.00015014 | 0.41390943 | ASV |
| **ASV304** | -0.007485 | 0.0040983 | -1.8234424 | 0.06823656 | -0.0158752 | 0.00032568 | 0.41890742 | ASV |
| **ASV48** | 0.00832114 | 0.00457403 | 1.81413425 | 0.06965706 | -0.000547 | 0.01753967 | 0.41918059 | ASV |
| **ASV712** | 0.02473058 | 0.01351562 | 1.82509585 | 0.06798657 | 0.00038864 | 0.05328596 | 0.42167381 | ASV |
| **ASV3** | -0.0084392 | 0.0046937 | -1.8060693 | 0.07090752 | -0.0177429 | 0.00071669 | 0.42232695 | ASV |
| **ASV82** | 0.00409347 | 0.00235883 | 1.73717147 | 0.08235691 | -0.0004457 | 0.00879073 | 0.43639166 | ASV |
| **ASV546** | 0.0070009 | 0.0040281 | 1.74339053 | 0.08126542 | -0.0005657 | 0.01521558 | 0.4373007 | ASV |
| **ASV105** | -0.0046621 | 0.00266276 | -1.737861 | 0.0822354 | -0.0099213 | 0.00058212 | 0.43776906 | ASV |
| **ASV70** | -0.0037029 | 0.0021842 | -1.6989626 | 0.08932623 | -0.0079834 | 0.00056961 | 0.44037819 | ASV |
| **ASV58** | -0.0052103 | 0.0030212 | -1.719497 | 0.08552391 | -0.0111665 | 0.00073446 | 0.44759876 | ASV |
| **ASV27** | -0.0030617 | 0.00180674 | -1.6951229 | 0.0900521 | -0.0066297 | 0.00047086 | 0.45230821 | ASV |
| **ASV94** | -0.0024818 | 0.00144423 | -1.7194177 | 0.08553834 | -0.0053636 | 0.00034783 | 0.45508643 | ASV |
| **ASV387** | 0.00842096 | 0.00491848 | 1.70987069 | 0.08728979 | -0.0009811 | 0.0183066 | 0.4564837 | ASV |
| **ASV505** | 0.00857529 | 0.00497808 | 1.72707716 | 0.08415381 | -0.0004848 | 0.01926123 | 0.45974715 | ASV |
| **ASV97** | -0.0055875 | 0.00329174 | -1.7082802 | 0.08758437 | -0.0120579 | 0.00085097 | 0.46691715 | ASV |
| **ASV41** | 0.00556823 | 0.00334278 | 1.66886518 | 0.0951441 | -0.0008276 | 0.01230079 | 0.47959916 | ASV |
| **ASV14** | 0.00323065 | 0.00189647 | 1.70876205 | 0.08749504 | -0.0004098 | 0.00703063 | 0.48057585 | ASV |
| **ASV34** | -0.005072 | 0.00307179 | -1.6437825 | 0.10022114 | -0.0112212 | 0.00095401 | 0.49377663 | ASV |
| **ASV537** | 0.01101154 | 0.00687596 | 1.59552991 | 0.11059379 | -0.0006784 | 0.02702236 | 0.51913047 | ASV |
| **G: BIFIDOBACTERIUM** | -0.0071807 | 0.00219426 | -3.2879756 | 0.00100911 | -0.0115161 | -0.0029175 | 0.08477936 | Genus |
| **G: LACHNOSPIRACEAE FAMILY** | -0.0041852 | 0.00140062 | -2.9930506 | 0.00276204 | -0.0069458 | -0.0014554 | 0.12612225 | Genus |

| | | | | | | | | |
|---|---|---|---|---|---|---|---|---|
| **G: DESULFOVIBRIO** | -0.0161273 | 0.00609643 | -2.6548721 | 0.00793385 | -0.0287149 | -0.004609 | 0.17491611 | Genus |
| **G: COLIDEXTRIBACTER** | -0.0053179 | 0.00206327 | -2.5695621 | 0.01018272 | -0.0093954 | -0.0012846 | 0.18409546 | Genus |
| **G: FUSICATENIBACTER** | -0.0047185 | 0.00187526 | -2.5099011 | 0.0120765 | -0.0083984 | -0.0010337 | 0.19244421 | Genus |
| **G: LACHNOSPIRACEAE NK4A136 GROUP** | -0.0033387 | 0.00138403 | -2.4248631 | 0.01531417 | -0.0060438 | -0.0006309 | 0.20778439 | Genus |
| **G: BARNESIELLA** | -0.0048902 | 0.00205447 | -2.3798324 | 0.01732051 | -0.0089406 | -0.0008685 | 0.20876016 | Genus |
| **G: OSCILLOSPIRACEAE FAMILY** | -0.0056003 | 0.00236322 | -2.3828816 | 0.01717772 | -0.0102538 | -0.0009979 | 0.21397241 | Genus |
| **G: ENTEROCOCCUS** | -0.0065851 | 0.00280656 | -2.3423792 | 0.01916123 | -0.0123046 | -0.0011328 | 0.21639782 | Genus |
| **G: [EUBACTERIUM] HALLII GROUP** | -0.005858 | 0.00250375 | -2.3466231 | 0.01894441 | -0.010753 | -0.0009731 | 0.22079362 | Genus |
| **G: DIALISTER** | -0.0033084 | 0.00146604 | -2.2598637 | 0.02382971 | -0.0061907 | -0.0004413 | 0.23544579 | Genus |
| **G: UCG-005** | -0.0039882 | 0.001784 | -2.2415348 | 0.02499145 | -0.0075123 | -0.0005085 | 0.23801463 | Genus |
| **G: HAEMOPHILUS** | -0.00296 | 0.00133538 | -2.2220218 | 0.02628183 | -0.0055871 | -0.000337 | 0.24326442 | Genus |
| **G: BACTEROIDES** | -0.0038935 | 0.00180958 | -2.1490851 | 0.03162772 | -0.0074322 | -0.0003486 | 0.25948394 | Genus |
| **G: GORDONIBACTER** | -0.0046214 | 0.00226099 | -2.0451604 | 0.04083905 | -0.0090849 | -0.0002068 | 0.29429719 | Genus |
| **G: INTESTINIBACTER** | -0.0037523 | 0.00187568 | -2.0160076 | 0.04379918 | -0.0074237 | -0.0001014 | 0.29545764 | Genus |
| **G: ALISTIPES** | -0.0032307 | 0.00159488 | -2.0118054 | 0.04424048 | -0.0063482 | -8.266E-05 | 0.30066585 | Genus |
| **G: DTU089** | -0.0081764 | 0.00416069 | -1.9579339 | 0.05023778 | -0.0163634 | -8.363E-06 | 0.31629873 | Genus |
| **G: SUTTERELLA** | -0.006712 | 0.00347374 | -1.9295674 | 0.05366046 | -0.0136213 | 9.7326E-05 | 0.3255157 | Genus |
| **G: ERYSIPELOTRICHACEAE UCG-003** | -0.0039743 | 0.00207952 | -1.9157725 | 0.05539406 | -0.0080495 | 9.7619E-05 | 0.33362824 | Genus |
| **G: [RUMINOCOCCUS] TORQUES GROUP** | -0.0038578 | 0.0020784 | -1.8570125 | 0.06330937 | -0.0079396 | 0.00021959 | 0.35066156 | Genus |
| **G: CANDIDATUS SOLEAFERREA** | -0.0036372 | 0.0019608 | -1.8442698 | 0.06514383 | -0.0075125 | 0.00020606 | 0.35370709 | Genus |
| **G: AKKERMANSIA** | -0.0025114 | 0.00137248 | -1.833876 | 0.06667244 | -0.0052102 | 0.0001741 | 0.36005771 | Genus |
| **G: PORPHYROMONAS** | -0.0092352 | 0.00511732 | -1.8119502 | 0.06999389 | -0.0195197 | 0.00064047 | 0.36644401 | Genus |
| **G: CLOSTRIDIUM SENSU STRICTO 1** | -0.0022775 | 0.00126212 | -1.8101247 | 0.07027645 | -0.0047486 | 0.0001971 | 0.36808989 | Genus |
| **G: GEMELLA** | -0.0067439 | 0.00374418 | -1.7988242 | 0.0720465 | -0.0141543 | 0.00057944 | 0.37113308 | Genus |
| **G: PEPTOCOCCACEAE FAMILY** | -0.0117762 | 0.00670309 | -1.7465513 | 0.08071575 | -0.025027 | 0.00137363 | 0.38045805 | Genus |
| **G: PREVOTELLA** | -0.0053747 | 0.00304217 | -1.7659309 | 0.07740746 | -0.0115208 | 0.000602 | 0.38422879 | Genus |
| **G: ROSEBURIA** | -0.0024493 | 0.00145008 | -1.695314 | 0.09001587 | -0.0053007 | 0.0003933 | 0.41055067 | Genus |

| | | | | | | | | |
|---|---|---|---|---|---|---|---|---|
| **G: ANAEROFUSTIS** | -0.0062892 | 0.00371759 | -1.6966156 | 0.08976938 | -0.0136102 | 0.00096651 | 0.41399052 | Genus |
| **G: [RUMINOCOCCUS] GNAVUS GROUP** | -0.0029643 | 0.00180392 | -1.6463499 | 0.09969222 | -0.0065063 | 0.00058546 | 0.41686138 | Genus |
| **G: PARABACTEROIDES** | -0.003186 | 0.00191386 | -1.667061 | 0.09550229 | -0.0069586 | 0.00056614 | 0.41902758 | Genus |

*Supplementary Table 5 PERMANOVA output of ASD sub-types clustered on Euclidean distances using all datasets combined.*

| TERM | DEGREES OF FREEDOM | SUM OF SQUARES | $R^2$(%) | F-STATISTIC | P-VALUE | P-ADJUSTED VALUES (BENJAMINI-HOCHBERG) |
|---|---|---|---|---|---|---|
| **ASD.DSM.V** | 1 | 123746.743 | 1.0 | 1.260 | 0.069 | 1.000 |
| **ZYRTEC** | 1 | 97292.720 | 0.8 | 0.990 | 0.444 | 1.000 |
| **NEURODEVELOPMENT.DELAYS** | 1 | 92699.504 | 0.8 | 0.944 | 0.571 | 1.000 |
| **VITAMIN.SUPPLEMENTS** | 1 | 93050.275 | 0.8 | 0.947 | 0.666 | 1.000 |
| **NEUROLOGICAL.CONDITIONS** | 1 | 84651.867 | 0.7 | 0.862 | 0.755 | 1.000 |
| **ASD.DSM.IV** | 1 | 84095.244 | 0.7 | 0.856 | 0.881 | 1.000 |
| **PPD** | 1 | 75657.428 | 0.6 | 0.770 | 0.887 | 1.000 |
| **CLARITIN** | 1 | 74630.045 | 0.6 | 0.760 | 0.892 | 1.000 |
| **GENDER_FEMALE** | 1 | 82295.830 | 0.7 | 0.838 | 0.913 | 1.000 |
| **FISH.OIL** | 1 | 80361.291 | 0.7 | 0.818 | 0.920 | 1.000 |
| **ALBUTEROL** | 1 | 77289.744 | 0.7 | 0.787 | 0.930 | 1.000 |
| **MELATONIN** | 1 | 74025.508 | 0.6 | 0.754 | 0.932 | 1.000 |

| | | | | | | |
|---|---|---|---|---|---|---|
| **BEHAVIORAL.EMOTIONAL.DISORDERS** | 1 | 61383.184 | 0.5 | 0.625 | 0.961 | 1.000 |
| **PROBIOTIC** | 1 | 79350.809 | 0.7 | 0.808 | 0.962 | 1.000 |
| **ASD.ICD** | 1 | 80776.826 | 0.7 | 0.822 | 0.966 | 1.000 |
| **DIGESTIVE.ISSUES** | 1 | 70474.920 | 0.6 | 0.717 | 0.987 | 1.000 |
| **ASTHMA** | 1 | 65938.251 | 0.6 | 0.671 | 0.994 | 1.000 |
| **ALLERGIES** | 1 | 63962.438 | 0.5 | 0.651 | 1.000 | 1.000 |
| **ADHD** | 1 | 55778.602 | 0.5 | 0.568 | 1.000 | 1.000 |
| **RESIDUAL** | 105 | 10315417.540 | 87.2 | NA | NA | NA |
| **TOTAL** | 124 | 11832878.771 | 100.0 | NA | NA | NA |
